# Stoichiometric 14-3-3ζ binding promotes phospho-Tau microtubule dissociation and reduces aggregation and condensation

**DOI:** 10.1101/2024.03.15.585148

**Authors:** Janine Hochmair, Maxime C. M. van den Oetelaar, Leandre Ravatt, Lisa Diez, Lenne J. M. Lemmens, Renata Ponce-Lina, Rithika Sankar, Maximilian Franck, Gesa Nolte, Ekaterina Semenova, Satabdee Mohapatra, Christian Ottmann, Luc Brunsveld, Susanne Wegmann

## Abstract

The microtubule (MT) association of protein Tau is decreased upon phosphorylation. Increased levels of phosphorylated Tau in the cytosol pose the risk of pathological aggregation, as observed in neurodegenerative diseases. We show that binding of 14-3-3ζ enhances cytosolic Tau solubility by promoting phosphorylated Tau removal from MTs, while simultaneously inhibiting Tau aggregation both directly and indirectly via suppression of condensate formation. These 14-3-3ζ activities depend on site-specific binding of 14-3-3 to Tau phosphorylated at S214 and S324. At sub-stoichiometric 14-3-3ζ concentrations, or in the presence of other 14-3-3ζ binding partners, multivalent electrostatic interactions promote Tau:14-3-3ζ co-condensation, offering a phosphorylation-independent mode of Tau-14-3-3ζ interactions. Given the high abundance of 14-3-3 proteins in the brain, 14-3-3 binding could provide efficient multi-modal chaperoning activity for Tau in the healthy brain and be important for preventing Tau aggregation in disease.

## Introduction

The intrinsically disordered microtubule associated protein Tau is a highly soluble and abundant cytosolic protein. Tau is mainly expressed in neurons of the central nervous system where it reaches average cytosolic concentrations of about 2 μM^1–3^. In Alzheimer’s disease (AD), frontotemporal dementia (FTD), and related neurodegenerative brain diseases, Tau aggregates into insoluble amyloid-like fibrils that accumulate in the neuronal cytoplasm as neurofibrillary tangles (NFTs) and correlate with neurotoxicity, neuronal loss, and cognitive decline in these diseases^4,5^. Tau’s intrinsic propensity to self-interact and aggregate is mediated by short amino acid motifs in its C-terminal repeat domain (TauRD, containing four pseudo-repeats R1-R4;^6–8^). Interestingly, the TauRD also constitutes large parts of the microtubule (MT) binding region of Tau^9,10^, suggesting a competition between Tau aggregation and MT binding.

Phosphorylation in and around the MT binding domain reduces the affinity of Tau for the MT surface^11–13^ and seems to be necessary for Tau dissociation from the MT surface to enable MT dynamics^14^ and regulate motor protein transport^15^. In pathological conditions, phosphorylated Tau (phospho-Tau) accumulates in NFTs^5^, which led to the current working model in which phosphorylation-induced MT dissociation increases the concentration of phospho-Tau in the cytosol and thereby permits its aggregation. However, soluble phospho-Tau is abundant in the healthy human brain^16^ and enriched during neurodevelopment^17,18^, yet does not aggregate in these conditions. In fact, most - if not all - cellular Tau appears to be phosphorylated to some degree^11,19^, indicating that efficient molecular mechanisms exist to prevent phospho-Tau aggregation in the healthy brain.

Tau can be phosphorylated by a number of abundant kinases, with the majority of Tau’s >80 putative phosphorylation sites being located in and around TauRD^20,21^, suggesting that Tau phosphorylation is more than a signal for MT dissociation and a trigger of aggregation. For example, cAMP-dependent protein kinase A (PKA) phosphorylates Tau on multiple residues in and around TauRD (main sites: pS214/ pS324/ pS356/ pS409/ pS416;^22^) but spares major Tau phospho-epitopes related to Tau pathology^23,24^, suggesting that PKA phosphorylation may contribute to Tau dissociation from MTs but not to its aggregation. Notably, PKA phosphorylation can also prime Tau phosphorylation by other kinases, like GSK3β and Cdk5, which then leads to Tau being modified on pathology associated phospho-sites (e.g., pS202, pT205, pS396, and pS404;^25,26^).

Members of the 14-3-3 protein family are important hubs within protein-protein interaction (PPI) networks^27^ that typically bind their “client” proteins dependent on phosphorylation. Two specific phospho-serine/phospho-threonine sites in a client can jointly increase the client’s binding affinity to 14-3-3 dimers^28–31^. In human cells, seven 14-3-3 isoforms (β, γ, ε, η, σ, τ, and ζ) are expressed, each from a different gene. In the brain, 14-3-3 proteins constitute about 1% (w/w) of the soluble proteome^32,33^ and are crucially involved in neurodevelopment and synaptic health^34–40^. Furthermore, 14-3-3 proteins were reported to interact with and modulate the aggregation of proteins accumulating in neurodegenerative protein aggregation diseases, including alpha-synuclein in Parkinson’s disease^41^, GFAP relevant in Alexander disease^42^, as well as Tau in AD and FTD^43–45^. Previous immunohistology and co-immunoprecipitation (co-IP) experiments suggested that 14-3-3 proteins bind and/or co-aggregate with highly phosphorylated Tau in NFTs of *postmortem* AD brains^36,43,46,47^. These observations were widely interpreted as a pro-aggregation effect of 14-3-3 on Tau. *In vitro* Tau aggregation assays revealed that 14-3-3 triggers the aggregation of unphosphorylated Tau but in contrary decreases aggregation of Tau phosphorylated by PKA^48–50^.

Binding of different 14-3-3 isoforms to Tau was reported to depend on the presence of phosphate groups at PKA-target sites in Tau^48,49^, mainly residues S214 in the proline-rich domain (P1) and S324 in the third repeat (R3)^51,52^, which bind tightly in the binding groove of 14-3-3 dimers^44^. Because of the high 14-3-3 abundance in the brain, binding of 14-3-3 to Tau phosphorylated at pS214/pS324 (Tau_pS214/pS324_) could provide a robust mechanism for promoting the solubility of PKA-phosphorylated Tau (PKA-Tau) and preventing its aggregation. In turn, deregulation of 14-3-3:Tau interactions could allow for PKA-Tau aggregation in neurodegenerative diseases. Notably, activating cAMP pathways – and therefore PKA – was shown to be protective against Tau aggregation and toxicity in Tau transgenic mice^53^, whereas a reduced abundance of 14-3-3 proteins in AD brains^46,54^ may indirectly promote the intra-neuronal aggregation of highly phosphorylated Tau typical for the disease.

Here, we show that pS214 and pS324 phosphorylation-dependent binding of 14-3-3 increases Tau solubility and decreases Tau aggregation in neurons. Molecularly, these effects are based on (i) 14-3-3 mediated “scavenging” of Tau_pS214/pS324_ molecules from the MT surface based on Tau binding competition between MTs and 14-3-3, and (ii) the inhibition of Tau_pS214/pS324_ condensation, potentially *en-route* to the formation of pathological Tau species^3,55^, and amyloid-like aggregation by stoichiometric 14-3-3 binding. However, sub-stochiometric 14-3-3 concentrations promote Tau condensation and may therefore increase the risk for Tau aggregation. Together, our data elucidate the importance of 14-3-3 proteins in modulating Tau pathobiology, in which the formation of stoichiometric, soluble 14-3-3:Tau_pS214/pS324_ complexes reduces the risk of un-chaperoned phospho-Tau accumulation and spontaneous aggregation in the cytosol at the earliest step.

## Results

### 14-3-3 binding suppresses Tau aggregation in neurons

In the healthy brain, a robust occupation of soluble (not bound to MTs) phospho-Tau molecules by 14-3-3 and other chaperoning proteins could be essential for efficient prevention of Tau aggregation. In contrast, inhibition of chaperone binding could enable or enhance phospho-Tau aggregation.

To test whether the binding of 14-3-3 proteins can suppress neuronal Tau aggregation, we expressed pro-aggregant FTD-mutant human Tau (eGFP-Tau^P301L/S320F^) in primary hippocampal mouse neurons, which leads to spontaneous eGFP-Tau^P301L/S320F^ aggregation into NFT-like, fibrillar aggregates in a subset of neurons (∼2% NFTs in DMSO control; Fig. 1a,b). Co-immunoprecipitation from neuronal lysates confirmed the association of 14-3-3 with Tau (Supplemental Fig. S1a). Treatment with BV02, an inhibitor of 14-3-3 proteins binding to phosphorylated clients^56,57^, increased the fraction of neurons with eGFP-Tau^P301L/S320F^ aggregates in a dose-dependent manner (∼4% NFTs at 10 μM BV02, ∼7% NFTs at 40 μM BV02). In contrast, increasing 14-3-3 binding by treatment with fusicoccin (50 μM;^58,59^) decreased neuronal Tau aggregation (∼1% NFTs) and simultaneously increased the concentration (=fluorescence intensity) of soluble eGFP-Tau^P301L/S320F^ in the soma of neurons without aggregates (Fig. 1c). Together, these data suggested that 14-3-3 binding increases cytosolic Tau solubility, whereas reducing 14-3-3 binding promotes Tau aggregation in neurons.

**Figure 1.**
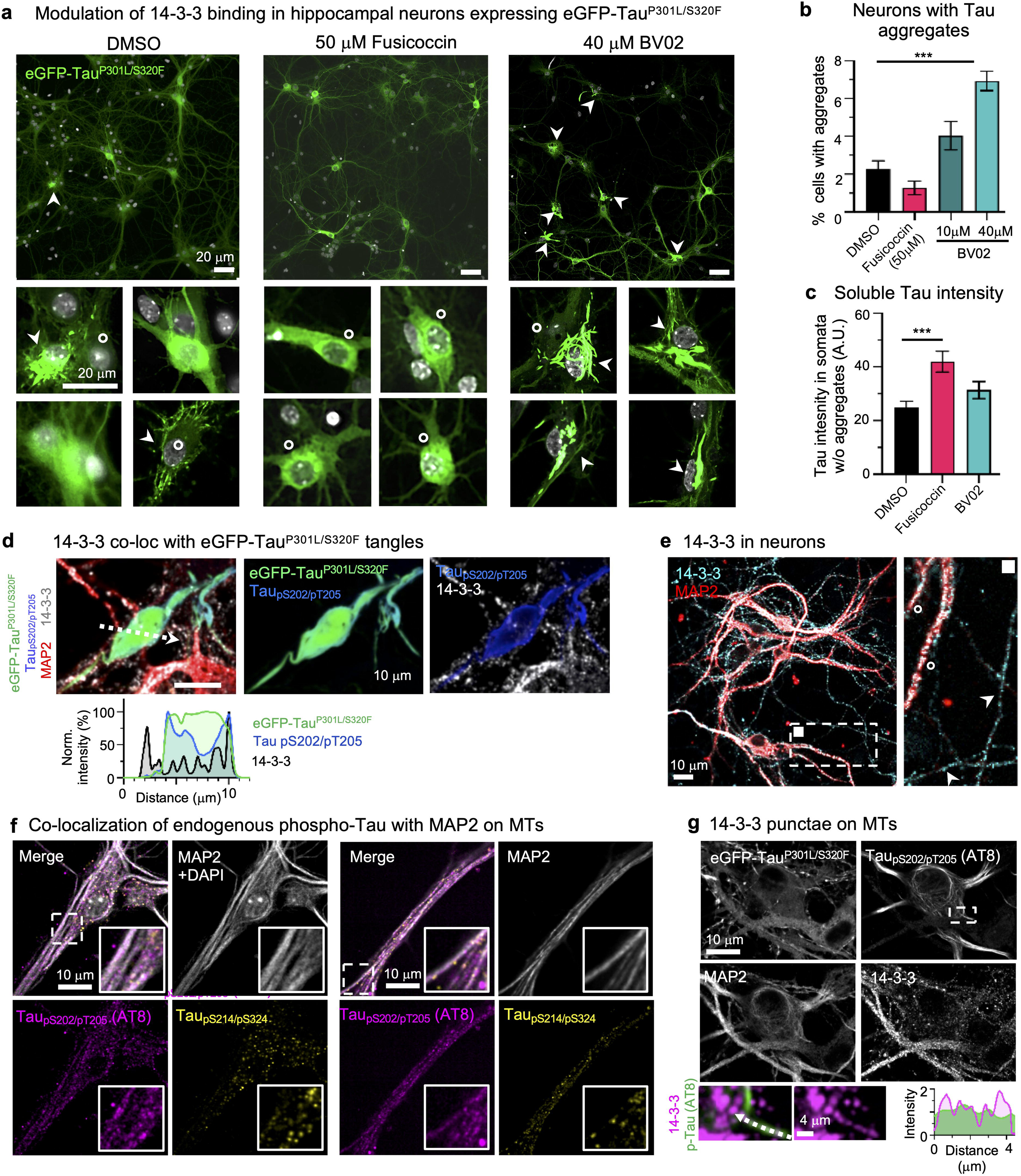
14-3-3ζ binding inhibits neuronal Tau aggregation. **a,** Representative images of primary hippocampal neurons (DIV12) transduced with AAV to express aggregating FTD-mutant eGFP-Tau^P301L/S320^. Neurons were treated with DMSO (control), fusicoccin (50 μM), or BV02 (40 μM) for 48h, starting 3 days after AAV transduction. Zoom-ins show examples of cell bodies with fibrillar Tau aggregates (white arrow heads) or high soluble Tau (white circles). Scale bars = 20 μm. **b,** Quantification of Tau aggregates in neurons upon treatment with DMSO, fusicoccin (50 μM), or BV02 (10 or 40 μM). Data shown as mean±SEM, N=4 analyzed images, one-way ANOVA with Tukey post-test. **c,** Quantification of soluble eGFP-Tau (mean intensity) in the soma of neurons without aggregates. Data shown as mean±SEM, N=4 analyzed images, one-way ANOVA with Tukey post-test. **d,** Representative image of fibrillar eGFP-Tau aggregate in neurons immunolabeled for MAP2 (red), 14-3-3 (white), and Tau_pS202/pT205_ (blue; AT8 antibody). Line plot along white arrow shows no enrichment and limited colocalization of granular 14-3-3 in tangle-like Tau aggregate. Scale bar = 10 μm. **e,** Representative image of neurons immunolabeled for 14-3-3 (cyan) and MAP2 (red). Zoom-in shows granular labeling of 14-3-3 in MAP2+ dendrites (white circles) and MAP2-axons (white arrow heads). Principle dendrites appear rich in 14-3-3. Scale bar = 10 μm. **f,** Neurons immunolabeled for MAP2 and phospho-Tau variants indicate stronger colocalization of endogenous Tau_pS202/pT205_ (AT8 epitope) than Tau_pS214/pS324_ with MAP2+ microtubules (MTs). Scale bars = 10 μm. **g,** Example fluorescence image of a neuron expressing eGFP-Tau^P301L/S320F^, with somatodendritic MAP2-coated microtubules that are also coated with Tau_pS202/pT205_ (AT8 antibody). Zoom-ins show that 14-3-3 granules (pink) align with Tau_pS202/pT205_ (green) coated microtubules (compare line plot along white arrow). Scale bars = 10 μm, 4 μm for zoom-in.

To test whether 14-3-3 would colocalize with aggregated or soluble phospho-Tau, we performed immunostainings on eGFP-Tau^P301L/S320F^ expressing neurons. We found that, in contrast to previous reports from human brain immunostainings^43,46,47^, 14-3-3 was mostly excluded from NFT-like eGFP-Tau^P301L/S320F^ aggregates in our neuron model (Fig. 1d), in which 14-3-3 showed a punctate (=granular) staining pattern in dendrites, axons, and somata (Fig. 1e), similar to previous reports from human brain^47^ without Tau NFTs.

Interestingly, Tau phosphorylated at PKA-target sites pS214 and pS324, two of the phosphorylation sites mainly relevant for 14-3-3:Tau binding^52^, showed limited colocalization with MTs and a cytoplasmic, granular distribution, similar to 14-3-3 (Fig. 1f). In contrast, a Tau phospho-epitope previously not reported as relevant for 14-3-3:Tau binding (pS202/pT205, target sites of, e.g., Cdk5 and GSK3β) stained microtubules in naïve and eGFP-Tau^P301L/S320F^ overexpressing neurons (Fig. 1f). 14-3-3 granules seemed to arrange along Tau_pS202/pT205_ coated MTs (Fig. 1g). These data indicate that Tau_pS214_ binds MTs less efficiently than Tau_pS202/pT205_, which could be related to its 14-3-3 binding.

### Efficient 14-3-3ζ binding depends on Tau phosphorylation at residues S214 and S324

Given the previously suggested link between 14-3-3, Tau and MT stability in axon development and regeneration^35,60^, we established an *in vitro* model for the phosphorylation-dependent binding of Tau to 14-3-3, in order to probe the effect on Tau’s MT binding.

We generated phospho-null mutants and phospho-variants (by *in vitro* phosphorylation with purified kinases) of recombinant full-length human Tau (2N4R isoform; Fig. 2a) and determined their binding affinities to the 14-3-3ζ isoform, which is the most abundant isoform in the brain^61^.

**Figure 2.**
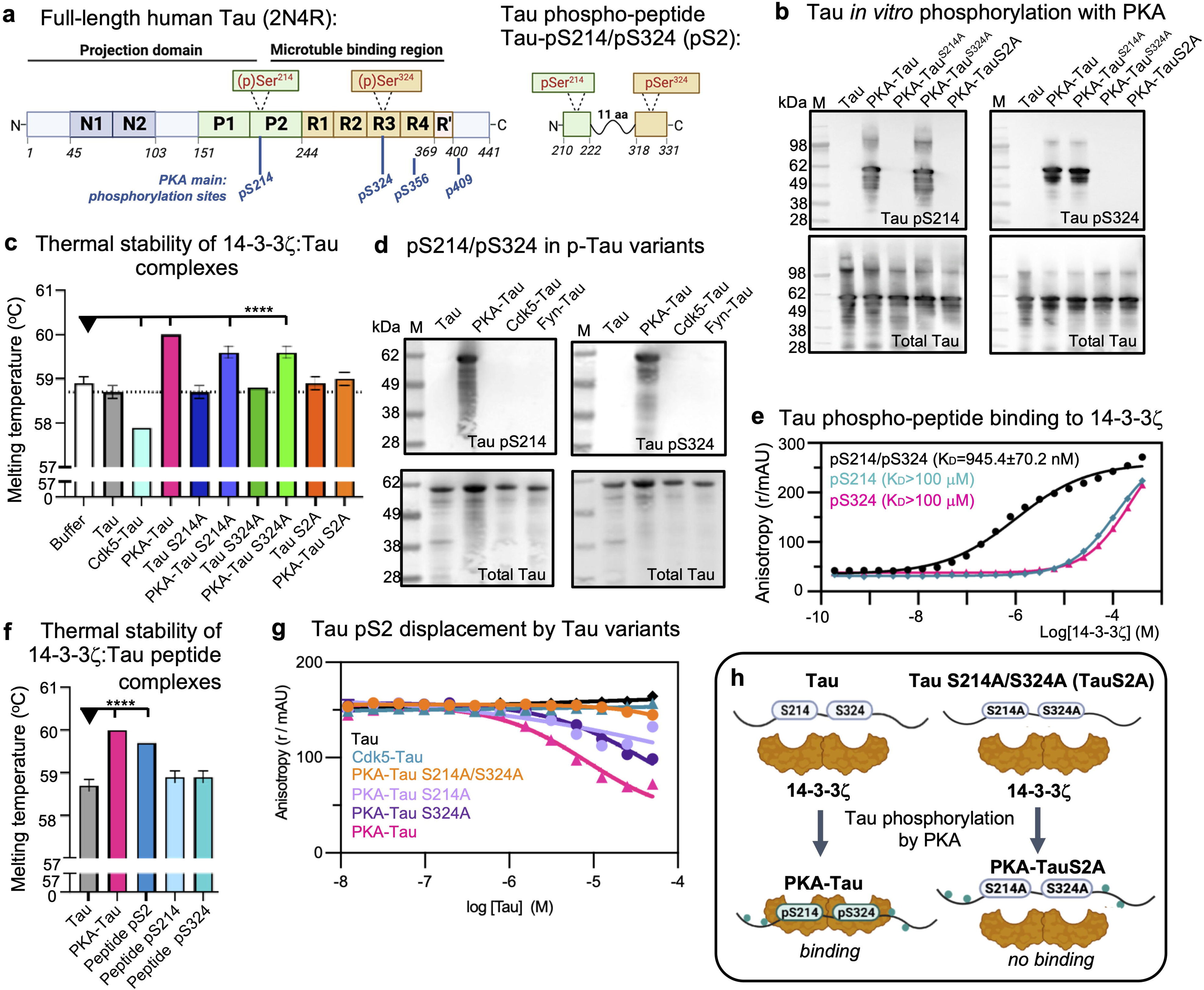
14-3-3ζ binding to Tau depends on phosphorylation at S214 and S324. **a,** Left: Domain structure of the longest human Tau isoform (2N4R, 441 aa) consisting of the N-terminal projection domain with two N-terminal inserts (N1, N2), the proline-rich domain (P1+P2), and the C-terminal microtubule binding region that includes four pseudo-repeats (R1-R4) and short sequences up- and downstream of these. Positions of phosphorylation sites on serine residues Ser214 in P2 and Ser324 in R3 are indicated. Right: Domain structure of phospho-peptide Tau_pS214/pS324_ (pS2) having 38 amino acids (Tau^210-222(pS214)^-GGGSGGGSGGG-Tau^318-331(pS324)^). **b,** Western blots of recombinant Tau variants *in vitro* phosphorylated using PKA kinase (Tau variants: wildtype Tau, Tau^S214A^, Tau^S324A^, Tau^S214A/S324A^ (Tau S2A)) using antibodies specific for Tau phospho-residues (Tau_pS214_ and Tau_pS324_) and total Tau. **c,** Thermal stability of 14-3-3ζ mixed with full-length Tau variants in their PKA-phosphorylated and non-phosphorylated forms. The higher the binding affinity, the higher is the temperature needed to melt Tau:14-3-3ζ complexes. PKA-phosphorylation increases binding of Tau to 14-3-3ζ, which is reduced upon S>A mutations in Tau^S214A^ and Tau^S324A^. Mutation of both serine residues abolishes Tau^S214A/S324A^ (=TauS2A) binding. Data shown as mean±SD, N=3 independent experiments. One-way ANOVA with Tukey post-test. Significance compared to “Buffer” (=14-3-3ζ alone) is indicated. **d,** Western blot of Tau_pS214_ and Tau_pS324_ sites in full-length Tau *in vitro* phosphorylated by different kinases. **e,** Representative fluorescent anisotropy measurements for the binding of Tau phospho-peptides pS2, Tau_pS214_, and Tau_pS324_ to 14-3-3ζ. 14-3-3ζ was titrated into 10 nM of respective fluorescein-labeled Tau peptide. Data shown as mean±SD, representative experiment with N=3 technical replicates (two more experiments shown in Supplemental Fig. S1c). **i,** Thermal stability of 14-3-3ζ mixed with Tau phospho-peptides (pS2, Tau_pS214_, and Tau_pS324_) compared to full-length Tau and PKA-Tau. Data shown as mean±SD, N=3 independent experiments. One-way ANOVA with Tukey post-test. Significance compared to “Tau” is indicated. **k,** Representative fluorescent anisotropy measurements for the competition between full-length phospho-Tau variants and pS2-FITC (10 nM) for the binding to 14-3-3ζ (1 μM). Data shown as mean±SD, N=3 technical replicates (two experiments shown in Supplemental Fig. S1d). **h,** Model: Full-length Tau binding to 14-3-3ζ depends on phosphorylation at S214 and S324.

The PKA target sites, S214 and S324^25^, were shown to be necessary for Tau binding to the 14-3-3σ isoform^44,60^. Our phospho-null mutants therefore contained serine-to-alanine mutations at S214 and S324 – individually or together (Tau^S214A^, Tau^S324A^, and Tau^S214A/S324A^ = TauS2A). *In vitro* phosphorylation with PKA introduced phosphates at S214 and S324 in wildtype Tau (PKA-Tau) but not at respective mutated serine residues in Tau^S214A^, Tau^S324A^, and Tau^S214A/S324A^, as confirmed by Western blot (Fig. 2b).

Next, we determined the binding strength between 14-3-3ζ and different PKA-Tau variants using thermal stability assays (measuring the temperature needed to denature 14-3-3:Tau complexes; Fig. 2c). 14-3-3ζ:PKA-Tau complexes demonstrated the highest thermal stability, followed by 14-3-3ζ:PKA-Tau^S214A^ and 14-3-3ζ:PKA-Tau^S324A^. By contrast, 14-3-3ζ:PKA-Tau^S214A/S324A^ was no more stable than 14-3-3:non-phosphorylated Tau. Furthermore, phosphorylation of Tau by Cdk5, which phosphorylates S202/T205 (Supplemental Fig. S1b;^55^) but not S214/S324 (Fig. 2d), also did not stabilize 14-3-3ζ:Tau interactions, and in fact decreased stability (Fig. 2c). This confirmed previous observations that, among all PKA phosphorylation sites in Tau, phosphorylation of S214 and S324 is individually and synergistically important for 14-3-3ζ:Tau binding.

To further assess whether the differences in thermal stability arose from different 14-3-3ζ:Tau binding affinities, we performed Tau phospho-peptide (pS2) displacement assays. Here, we titrated full-length Tau variants into pS2:14-3-3ζ and measured the reduction in fluorescently-labeled pS2 anisotropy resulting from its displacement from the complex. The pS2 Tau phospho-peptide consisted of two short Tau sequences with phosphate groups at S214 and S324, connected by an unstructured linker (pS2: 38 aa; Tau^210-222(pS214)^-GGGSGGGSGGG-Tau^318-331(pS324)^; Fig 2a; Supplemental Fig. S1c). pS2 showed efficient binding to 14-3-3ζ (K_D_∼1 μM), which also synergistically depended on phosphorylation at S214 and S324, as determined by fluorescent anisotropy and thermal stability assays, (Fig. 2e, f; Supplemental Fig. S1d). In the performed displacement assays (Fig. 2g, Supplemental Fig. S1e), pS2 was efficiently out-competed by full-length PKA-Tau, to a lesser degree by PKA-Tau^S214A^ and PKA-Tau^S324A^, but not by PKA-Tau^S214A/S324A^ or non-phosphorylated Tau. Cdk5-Tau was also not able to displace pS2 from 14-3-3ζ.

We note that PKA-Tau contains multiple phosphorylated residues in addition to pS214 and pS324^55,62,63^, as demonstrated in phos-tag gels by a pronounced upshift compared to non-phosphorylated Tau (Supplemental Fig. S1f). These additional phosphorylation sites should also be present in non-binding PKA-Tau^S214A/S324A^. Cdk5-Tau also contained phosphorylation at multiple sites (upshift in phos-tag gel), but not at pS214 and pS324 (Fig. 2d), and did not bind 14-3-3ζ. Therefore, additional Tau phosphorylation sites other than pS214 and pS324 may have less impact on Tau’s binding to 14-3-3ζ.

Together these data confirm the previously reported necessity of Tau phospho-epitopes pS214 and pS324 for the binding to 14-3-3^44,51,52^ and the relevance of PKA activity in this process, here for the 14-3-3ζ isoform (Fig. 2h). Phospho-epitopes pS214 and pS324 are located in regions relevant for Tau aggregation^64,65^ and microtubule binding ^10,66^; however, these specific phospho-sites have not been associated with Tau pathology in disease^23^. We thus suggest that 14-3-3 proteins bind physiological soluble rather than pathological aggregated Tau_pS214/pS324_ molecules and thereby promote their solubility.

### 14-3-3ζ binding enables dissociation of PKA-Tau from microtubules

Non-phosphorylated Tau has a high affinity to MTs (Tau:MTs K_D_ ≍ 1 μM;^9,67^) but the binding affinity drops upon phosphorylation of serine and threonine residues in and around TauRD. For example, PKA-Tau binds MTs with K_D_ ≍ 10 μM^68^. Since we found that 14-3-3ζ has a 10-fold stronger affinity for PKA-Tau (K_D_ ≍1 μM; Fig. 2f), we hypothesized that 14-3-3 would compete for PKA-Tau binding with the MT surface. Sequestration of PKA-Tau from MTs by 14-3-3ζ could explain why Tau_pS214_ staining was absent from MTs in neurons (Fig. 1f).

To test this idea, we performed *in vitro* MTs pelleting assays, in which MTs were polymerized in the presence of PKA-Tau or unphosphorylated Tau and in the absence or presence of 14-3-3ζ. After centrifugation, the pellet (P) fraction contained MTs and MT-bound Tau, whereas the supernatant (S) contained unbound Tau and 14-3-3. Indeed, the fraction of unbound PKA-Tau (increased S/P ratio) was significantly higher in the presence, compared to the absence, of 14-3-3ζ (Fig. 3a,b). The fraction of unbound non-phosphorylated Tau was unaffected by the presence of 14-3-3ζ, as expected in absence of 14-3-3ζ:Tau binding. These results thus indeed suggest that 14-3-3 reduces MT binding for PKA-Tau but not Tau (Fig. 3c).

**Figure 3.**
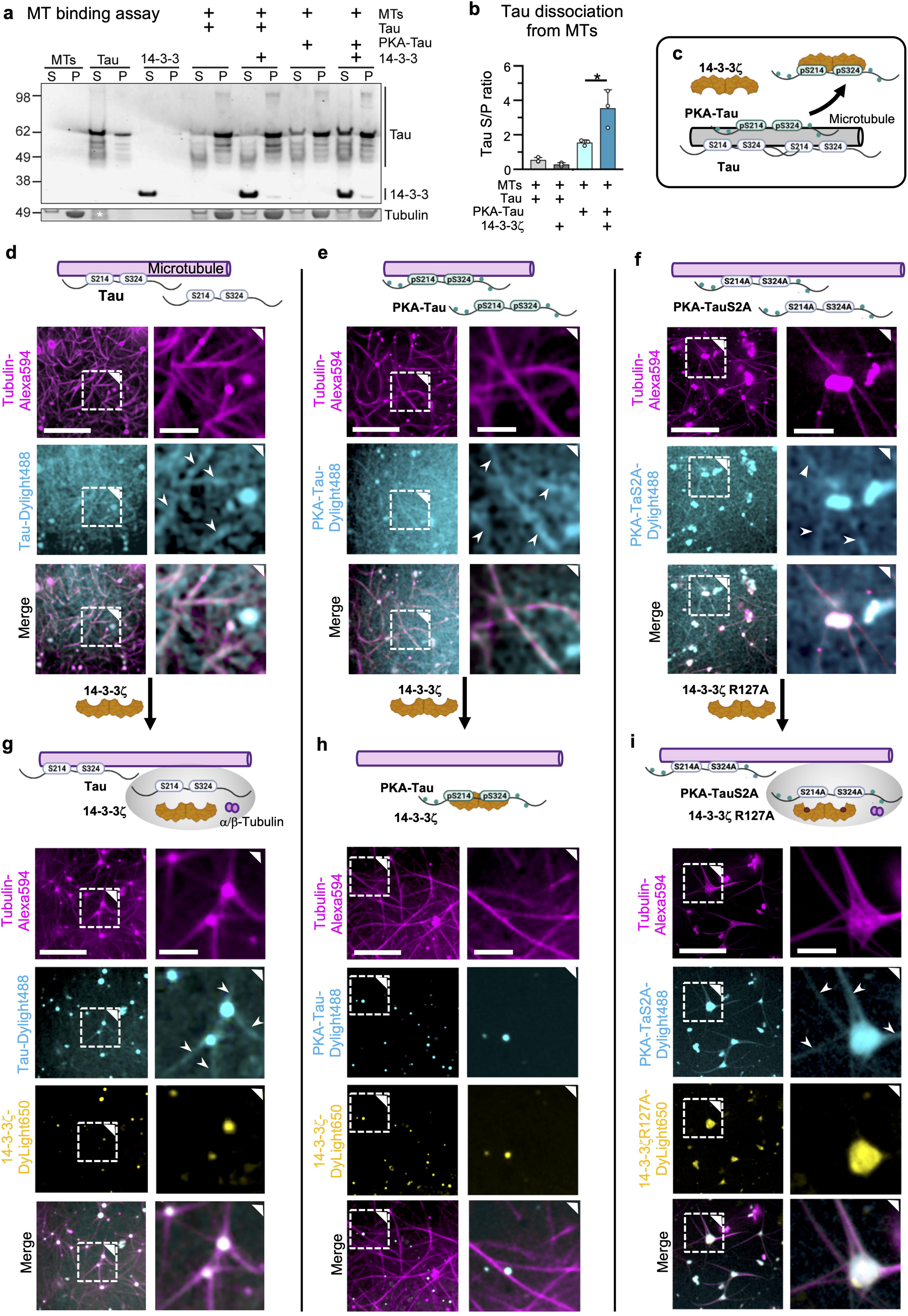
14-3-3ζ binding promotes Tau pS214/pS324 dissociation from microtubules. **a,** Western blot of MT binding assay for Tau and PKA-Tau with and without 14-3-3ζ. **b,** Quantification of MT binding assay. Supernatant (soluble unbound) to pellet (MT bound) S/P ratios for Tau and PKA-Tau are plotted. Data shown as mean±SD, N=3 independent assays, One-way ANOVA with Tukey post-test. **c,** Model for 14-3-3ζ promoting phospho-Tau detachment from MTs. **d,e,f,** Confocal images of *in vitro* MT formation with Tau variants (25 μM, 5% PEG) show Tau coating of MTs (white arrow heads) and remaining soluble Tau in solution. Non-phosphorylated Tau **(d)**, PKA-Tau **(e)**, and PKA-Tau^S2A^ **(f)**. Scale bars = 20 μm, and 5 μm in zoomed insets. **g,h,i,** Images of MT formation in the presence of both Tau (25 μM, 5% PEG) and 14-3-3ζ (12.5 μM). When 14-3-3ζ does not bind Tau (Tau+14-3-3ζ **(g)** and PKA-Tau^S2A^+14-3-3ζ^R127A^ **(i)**), Tau coats MTs (white arrow heads), no free Tau is in solution, and MTs grow from condensates containing Tau, 14-3-3ζ, and tubulin (=MT asters). When 14-3-3ζ is binding Tau (PKA-Tau+14-3-3ζ **(h)**), phospho-Tau is absent on MTs and few Tau:14-3-3ζ condensates form in solution, not attached to MTs. Scale bars = 20 μm, 5 μm in zoomed insets.

Intrigued by these findings, we aimed at visualizing the effect of 14-3-3ζ on the binding of PKA-Tau to MT *in vitro* by confocal microscopy using fluorescently labeled Tau and 14-3-3ζ variants (1-2% labeled proteins)^55,69^. We first polymerized MTs from tubulin-Alexa594 in the presence of non-phosphorylated Tau, PKA-Tau, or PKA-Tau^S214A/S324A^ (1% fluorescently Dylight488 labeled proteins). Surprisingly, we observed colocalization of all three Tau variants with MTs (Fig. 3d-f), indicating that PKA-phosphorylation alone was not sufficient to detach Tau from MTs, despite the decrease in MT binding affinity for PKA-Tau (K_D_ ≍ 10 μM^68^) compared to non-phosphorylated Tau (K_D_ ≍ 1 μM;^9,67^). In fact, the results from the MT pelleting assay also suggested that >50% (S/N ratio >1) of PKA-Tau was still bound to MTs (Fig. 3b).

When performing MT imaging in the presence of 14-3-3ζ, however, non-phosphorylated Tau (not binding 14-3-3ζ) still co-localized with MTs, whereas PKA-Tau (binding 14-3-3ζ) was now absent from MTs (Fig. 3g,h). These observations supported our hypothesis that 14-3-3ζ binding promotes the dissociation of PKA-Tau from MTs.

To prove that this was due to phospho-site specific binding of 14-3-3ζ, and not general electrostatic of unspecific interactions, we used a combination of non-binding mutants: PKA-Tau^S214A/S324A^, which contained PKA-induced phosphorylation except at pS214 and pS324, and 14-3-3ζ^R127A^, which contained an arginine-to-alanine mutation in its binding pocket and therefore binds phospho-Tau with lower affinity. In this non-binding combination, PKA-Tau^S214A/S324A^ remained on MTs (Fig. 3i), confirming that the binding of 14-3-3ζ was needed to sequester PKA-Tau from MTs. These findings may explain how overexpression of 14-3-3 can lead to a destabilization of the neuronal MT cytoskeleton^60^, i.e., by excessive sequestration of phospho-Tau from MTs.

### 14-3-3ζ co-condenses with non-binding Tau on microtubules

When imaging MTs, we noticed that all Tau variants formed condensates with free tubulin that attached to polymerized MTs (Fig. 3d-f), likely formed from excess, unbound Tau which recruits free tubulin. We previously reported similar observations^55,69^. When adding 14-3-3ζ to PKA-Tau:MT preparations, only few and smaller PKA-Tau:14-3-3ζ condensates could be observed (Fig. 3g), some of which also contained tubulin. In contrast, adding 14-3-3ζ to either Tau:MT or PKA-Tau^S214A/S324A^:MT preparations (both not binding 14-3-3ζ; Fig. 3f,h) promoted Tau:tubulin condensation and led to the formation of MT “junctions” with large condensates in their center. 14-3-3ζ co-partitioned into these condensates and did not coat outgrowing MTs, indicating that, when not binding Tau, 14-3-3ζ had a higher affinity to Tau condensates than to Tau bound to MTs or MT themselves. These observations were reminiscent of what we saw in neurons, where 14-3-3 granules partially aligned with MTs (Fig. 1g).

Together, the data suggest that in the absence of specific binding via Tau phosphosites pS214 and pS324, 14-3-3 may promote Tau condensation without interfering in Tau’s MT binding. However, phosphorylation dependent binding to 14-3-3 removes PKA-Tau from MTs via binding competition but may not promote its condensation. Notably, both Tau MT binding^69–71^ as well as pathological Tau aggregation^3,55,72,73^ have previously been reported to involve Tau condensation. Modulation of Tau condensation by 14-3-3 binding may therefore underlie the previously suggested involvement of 14-3-3 in both these processes^48,50,60^.

### Stochiometric binding of 14-3-3ζ inhibits PKA-Tau condensation

To examine the mechanisms behind 14-3-3ζ modulation of Tau condensation, we tested its impact on condensate formation of different phospho-Tau variants. Again, we phosphorylated Tau *in vitro* using different kinases that do (PKA) or do not (Cdk5 and Fyn) introduce phospho-groups at the crucial 14-3-3ζ binding sites S214 and S324. Increasing concentrations of 14-3-3ζ progressively inhibited PKA-Tau condensation (measured as condensate surface coverage). An equimolar concentration of 14-3-3ζ dimers (2 molecules of 14-3-3ζ to 1 molecule of PKA-Tau fully abolished PKA-Tau condensation (Fig. 4a). Cdk5-Tau and Fyn-Tau condensation decreased by only 20% at these concentrations. This suggested that condensate inhibition was driven by specific binding of 14-3-3ζ to PKA-Tau.

**Figure 4.**
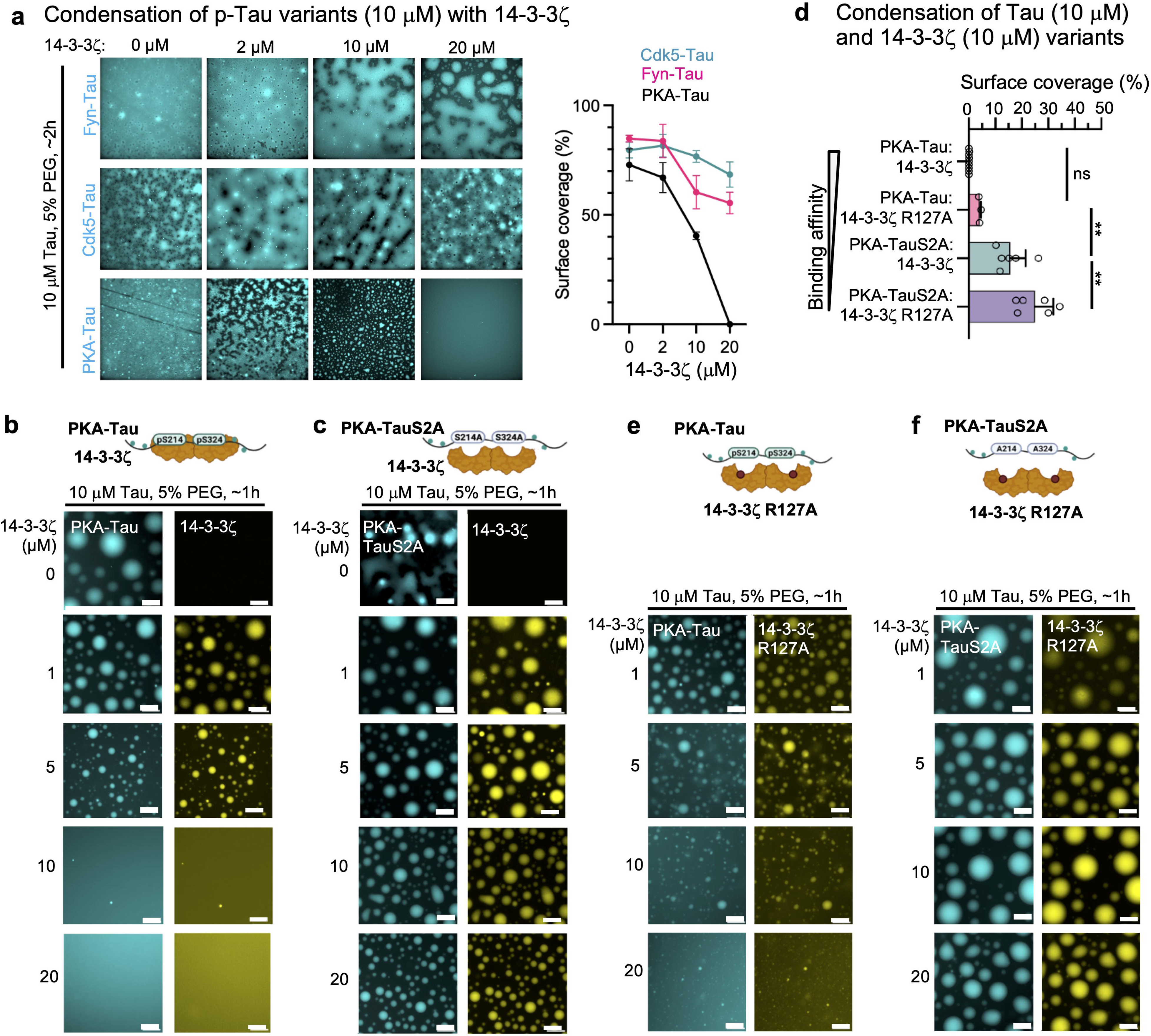
Stoichiometric binding of 14-3-3ζ inhibits Tau condensation. **a,** Representative images for the condensation of PKA-Tau, Cdk5-Tau and Fyn-Tau (each with 2% of phosphorylation matched Tau-DyLight488) at increasing 14-3-3ζ concentrations, recorded after 2-3 h (in 25 mM HEPES, pH 7.4, 10 mM NaCl, 1 mM DTT, 5% (w/v) PEG). Graph shows quantification of condensate surface coverage (%) for PKA-Tau, Cdk5-Tau and Fyn-Tau condensation. Data shown as mean±SD. N=3 images from 3 experimental replicates per condition. **b+c,** Representative images of 10 μM PKA-Tau (**a**), and PKA-TauS2A (**b**) with increasing 14-3-3ζ concentrations (0, 1, 5, 10, 20 μM; 2% Tau-DyLight488, 2% 14-3-3ζ-DyLight647). Scale bars = 20 μm. **d,** Quantification of condensate surface coverage (%) for 10 μM PKA-Tau and PKA-Tau^S214A/S324A^ (PKA-TauS2A) with 10 μM 14-3-3ζ or 14-3-3ζ^R127A^. Data shown as mean±SD. One-way ANOVA with Tukey post-test. Data points represent individual analyzed images, three images per condition, 2-3 experimental replicates. **e+f,** Representative images of 10 µM PKA-Tau (**c**), and PKA-TauS2A (**d**) with increasing 14-3-3ζ^R127A^ concentrations (1, 5, 10, 20 µM). Scale bars = 20 μm.

To test whether 14-3-3ζ binding to specifically pS214 and pS324 was necessary to inhibit Tau condensation, we incubated recombinant PKA-Tau (binding 14-3-3ζ) or PKA-Tau^S214A/S324A^ (not binding 14-3-3ζ) with increasing concentrations of 14-3-3ζ (10 μM Tau variants; 2% DyLight488-labeled PKA-Tau or PKA-Tau^S214A/S324A^; 2% DyLight650-labeled 14-3-3ζ) in condensation buffer (25 mM HEPES, pH7.4, ∼5 mM NaCl, 5% PEG). PKA-Tau and PKA-Tau^S214A/S324A^ each showed pronounced condensation in the absence of 14-3-3ζ (Fig. 4b,c), which confirmed previous observations^3,73^. Increasing concentrations of 14-3-3ζ inhibited condensation of PKA-Tau but not PKA-Tau^S214A/S324A^. 14-3-3ζ co-condensed with both Tau variants, but did not form condensates alone (Supplemental Fig. 2a). At equimolar concentrations of 14-3-3ζ (10 μM) and PKA-Tau (10 μM), when about half of PKA-Tau monomers are in complex with 14-3-3ζ dimers, no condensates were formed. In contrast, PKA-Tau^S214A/S324A^ condensation remained largely unaffected by 14-3-3ζ at this concentration ratio (Fig. 4d).

To further confirm the idea that Tau condensation can be tuned through 14-3-3 binding affinity, we modulated 14-3-3:PKA-Tau binding using 14-3-3ζ^R127A^ which reduces but does not abolish PKA-Tau binding. Incubating PKA-Tau with increasing concentrations of 14-3-3ζ^R127A^ led to inhibition of condensation, however, to a lesser degree than observed for wildtype 14-3-3ζ (Fig. 4e,f). This further supported a reciprocal relation between Tau condensation and 14-3-3ζ:Tau binding affinity. Condensation of PKA-Tau^S214A/S324A^ was not affected by 14-3-3ζ^R127A^. Tau condensation can thus be regulated via its 14-3-3 binding affinity.

In summary, these observations show that stoichiometric binding of 14-3-3ζ dimers to Tau monomers phosphorylated on S214 and S324 suppresses Tau condensation and therefore increases Tau solubility. This effect seemed to be largely independent of other Tau phosphorylation sites. Interestingly, suppression of biomolecular condensation by 14-3-3 binding has been suggested for other physiological and disease-associated 14-3-3 binding partners^74,75^.

### Tau:14-3-3ζ condensation depends on Tau availability and electrostatic interactions

Co-condensation of 14-3-3ζ with phosphorylated Tau was previously reported^76^, but most details of the molecular mechanisms and the connection to 14-3-3:Tau binding remain unclear. Our data showed that stoichiometric 14-3-3ζ binding suppresses PKA-Tau condensation in a concentration dependent manner. PKA-Tau bound to 14-3-3 appears to lose its ability to participate in condensate formation, and increasing 14-3-3 concentrations may deplete the pool of PKA-Tau available for condensation. We therefore hypothesized that re-establishing the availability of PKA-Tau should bring back PKA-Tau condensation.

To test this idea, we titrated the pS2 Tau peptide into an equimolar solution of PKA-Tau and 14-3-3ζ (10 μM each) that did not show condensation (Fig. 4b,d). At 10 μM pS2, PKA-Tau condensation indeed started to occur (Fig. 5a), suggesting that pS2 was outcompeting PKA-Tau from 14-3-3ζ dimers at this concentration, thereby re-enabling condensation of now un-bound PKA-Tau. Importantly, pS2 and 14-3-3ζ both co-enriched with PKA-Tau in condensates (Fig. 5b), indicating that 14-3-3ζ co-partitioned into PKA-Tau condensates while bound to pS2, similar to 14-3-3ζ:PKA-Tau complexes at lower 14-3-3ζ concentrations. Co-condensation of 14-3-3ζ with PKA-Tau therefore likely involved protein regions other than the Tau binding site of 14-3-3ζ.

**Figure 5.**
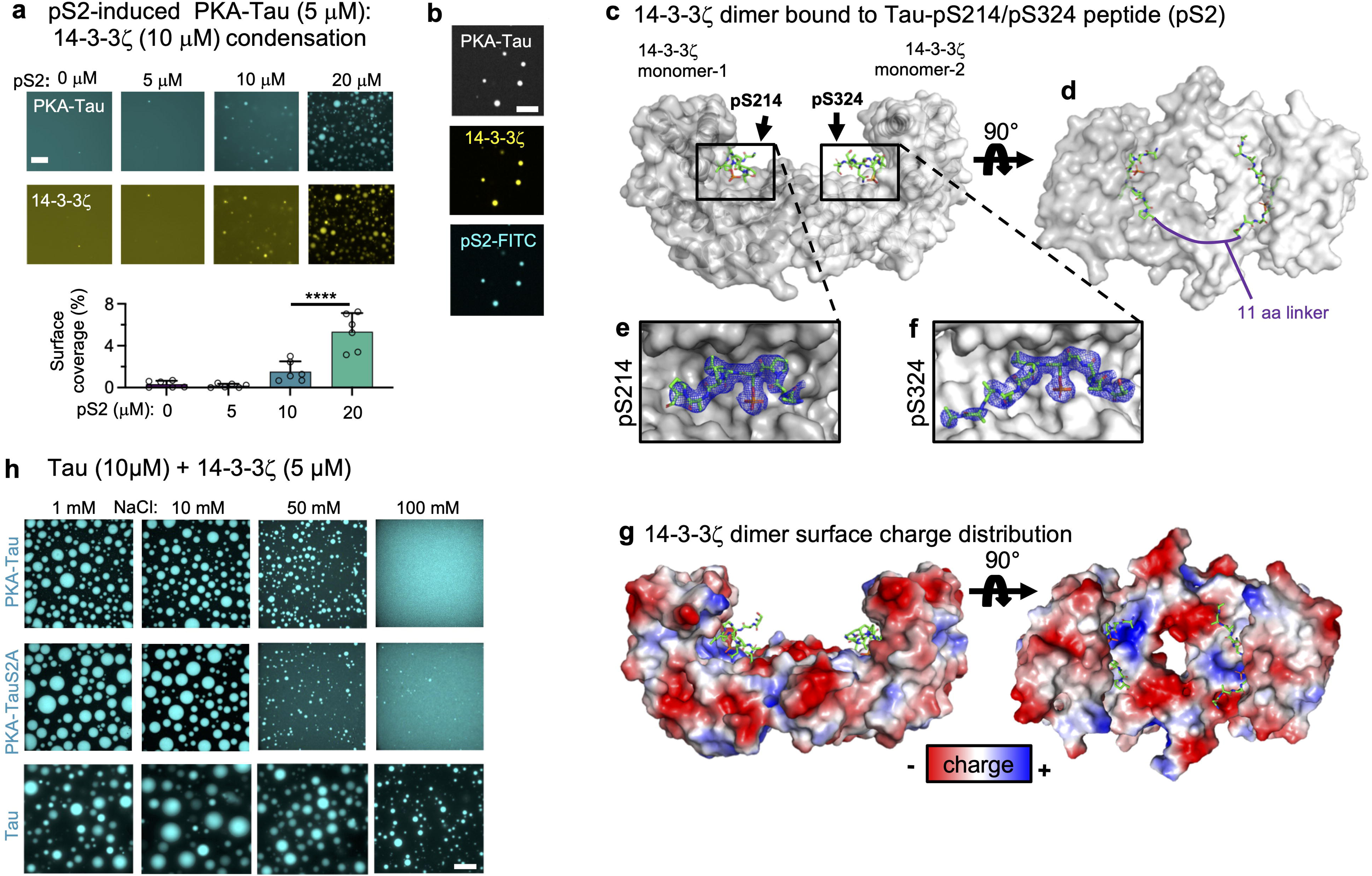
Tau:14-3-3ζ condensation depends on Tau availability and electrostatic interactions. **a,** Representative images of PKA-Tau (10 μM) with 14-3-3ζ (10 μM) and increasing concentrations of pS2 peptide (0, 5, 10, 20 μM). Scale bars = 20 μm. Quantification of condensate surface coverage (%) is shown in bar graph. Data shown as mean±SD. One-way ANOVA with Tukey post-test. Data points represent individual analyzed images, three images per condition, 3 experimental replicates. **b,** Representative images of condensates formed by PKA-Tau (10 μM, 2% PKA-Tau-DyLight405) with 14-3-3ζ (10 μM, 2% 14-3-3ζ-DyLight650) and pS2 (20 μM, 2% FITC-pS2). Scale bars = 10 μm. **c,** Crystal structure of 14-3-3ζ dimer (gray semi-transparent surface) in complex with pS214 and pS324 binding sites of pS2 (green rods). **d,** Top view of pS2 binding sites around pS214 and pS324 (green rods) in complex with 14-3-3ζ (gray semi-transparent surface). Solid purple line indicates connective unstructured 11 aa linker between the binding sites in pS2. **e+f,** Close-up of binding grooves in 14-3-3ζ monomers 1 and 2 (grey surface) in complex with pS2 binding sites (green rods). Final 2Fo-Fc electron density map of pS2 is shown as blue mesh (contoured at 1s). **g,** Surface charge distribution of 14-3-3ζ dimers mapped on crystal structure. Negatively charged areas are shown in red, positively charged ones in blue. Notably, the 14-3-3ζ binding pocket is positively charged, whereas much of the solvent exposed surface of 14-3-3ζ is negatively charged. **h,** Tau, PKA-Tau, and PKA-TauS2A (PKA-Tau^pS214A/pS324^) condensation with 14-3-3ζ with increasing NaCl concentrations (1, 10, 50, 100 mM). Scale bar = 10 μm.

The crystal structure of 14-3-3ζ dimers in complex with the pS2 peptide (Fig 5c,d; Supplemental Fig S3; Supplemental Table 1) revealed that pS2 bound in the standard 14-3-3ζ substrate binding groove established after 14-3-3 dimerization. pS2 residues pS214 and pS324 were binding to the two different 14-3-3ζ dimer subunits (Fig 5e,f), analogous to previously suggested binding modes of individual Tau^pS214^ and Tau^pS324^ peptides to the 14-3-3σ isoform^44^. The structure further showed many negatively charged areas on the 14-3-3ζ dimer surface (Fig. 5g). It is known from other studies that negatively charged biomolecules can co-condense with Tau^55,77,78^, driven by multivalent electrostatic interactions. Thus, co-condensation of Tau with 14-3-3ζ - being overall acidic (pI of 14-3-3ζ = 4-5;^31^) and having solvent-exposed negative charges on their surface (Fig. 5g) - could be driven by electrostatic interactions, e.g., with the positively charged TauRD. Indeed, recent NMR data showed that 14-3-3ζ, in addition to strong interactions with Tau phospho-sites pS214 and pS324, establishes many “weaker” interactions with the TauRD^22^, which may be involved in co-condensation.

We determined whether electrostatic interactions contribute to Tau:14-3-3ζ and PKA-Tau:14-3-3ζ condensation using different buffer salt (NaCl) concentrations that screen electrostatic protein-protein interactions to different degrees. Tau:14-3-3ζ condensates became smaller with increasing NaCl concentration, but remained observable at physiological salt concentrations (100 mM NaCl) (Fig. 5h). Electrostatic interactions thus played a role in stabilizing Tau:14-3-3ζ condensates. Additionally, we found that 14-3-3ζ promoted Tau condensation at net charge-matched concentrations (net charge of negative charges in 14-3-3ζ molecules = net charge of positive charges in Tau; 1-5 μM 14-3-3ζ at 10 μM Tau) (Supplemental Fig. S4), which is typical for electrostatically driven Tau condensation^78,79^.

For PKA-Tau:14-3-3ζ and PKA-Tau^S214A/S324A^:14-3-3ζ, however, 100 mM NaCl fully inhibited condensation. For the PKA-Tau variants, having a lower net charge because of negative charges added by phosphate groups, co-condensation with 14-3-3 may be more sensitive to buffer ion strength because of overall weaker multivalent electrostatic interactions with 14-3-3ζ. Thus, for phospho-Tau variants binding 14-3-3, both weaker multivalent interactions and masking of Tau stretches relevant for Tau condensation most likely contribute to inhibition of co-condensation.

### 14-3-3ζ binding prevents Tau amyloid aggregation but not condensate maturation

When monitored over time, Tau condensates exhibit a “maturation” process, during which Tau molecules lose their mobility in condensates and form species that seed Tau aggregation^3,55^. Independent of condensation, Tau aggregation can also be induced by direct aggregation into amyloid-like fibrils^79^. Binding of Tau by 14-3-3 seems to work efficiently against both Tau aggregation pathways, as demonstrated by the inhibition of PKA-Tau condensation *in vitro* (Fig. 4b) and Tau aggregation in neurons (Fig. 1a,b). However, the promotion of Tau condensation by 14-3-3 in non-binding conditions could induce Tau aggregation via condensate maturation. We investigated *in vitro* how 14-3-3 would modulate Tau amyloid aggregation and condensate maturation in phospho-site dependent binding and non-binding conditions.

First, to test whether Tau amyloid aggregation was modulated by 14-3-3ζ binding, we performed Thioflavin-T (ThioT) aggregation assays for Tau and PKA-Tau in the absence or presence of 14-3-3ζ. Here, we used the FTD-mutant pro-aggregant Tau^ΔK280^ to facilitate timely amyloid aggregation within 1-2 days. 14-3-3ζ inhibited PKA-Tau^ΔK280^ aggregation but promoted amyloid aggregation of non-phosphorylated Tau^ΔK280^ (Fig. 6a), similar to previous observations^48^. This indicated that occupation of TauRD by 14-3-3ζ in PKA-Tau (binding conditions) can prevent aggregation into amyloid-like fibrils, whereas 14-3-3 may act as polyanionic co-factor triggering Tau aggregation in non-binding conditions.

**Figure 6.**
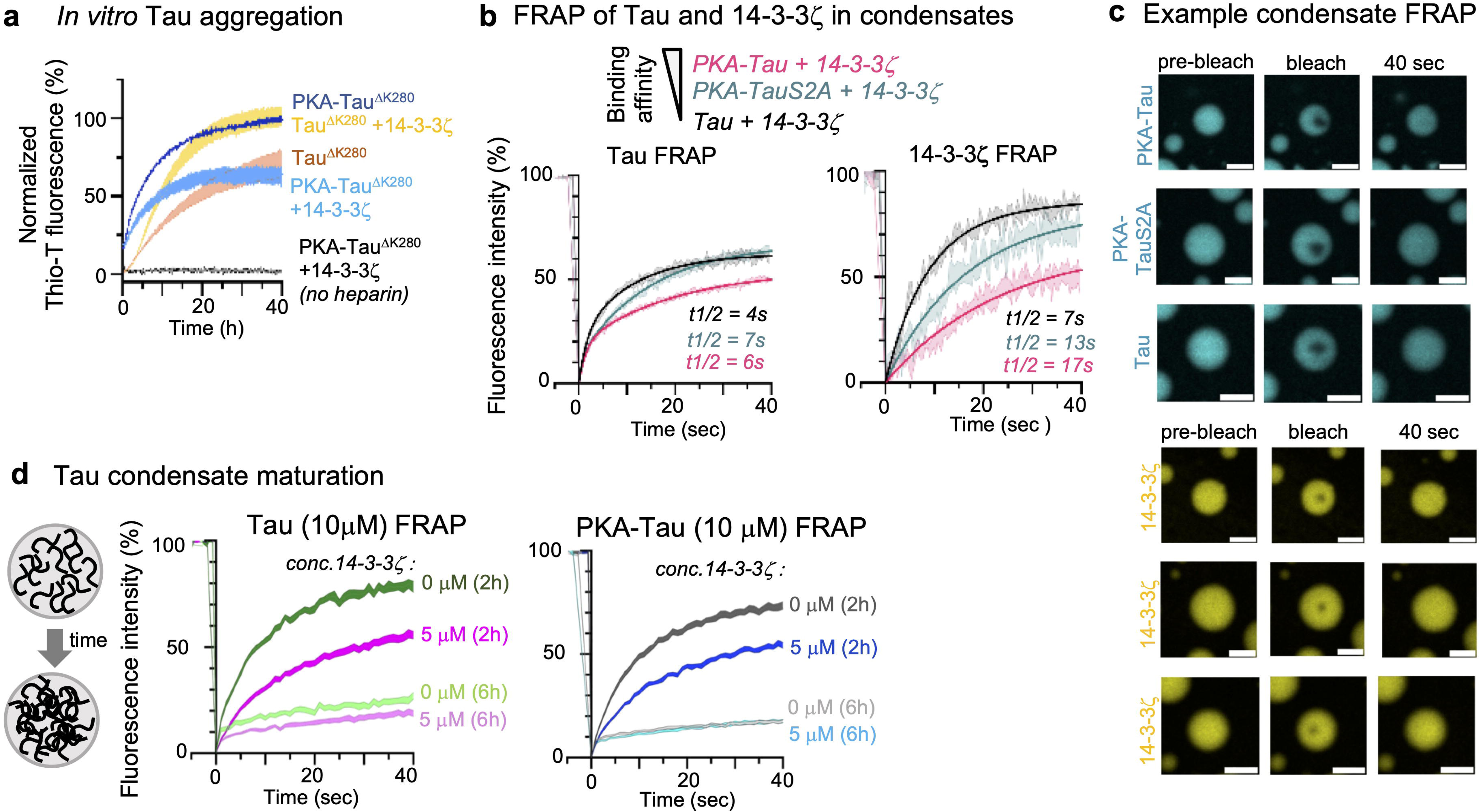
14-3-3ζ inhibits Tau amyloid aggregation but not condensate polymerization. **a,** Thioflavine-T (Thio-T) assay of pro-aggregant FTD-mutant Tau^ΔK280^ or PKA-Tau^ΔK280^ aggregation (triggered with heparin) in the absence and presence of 14-3-3ζ. N=3 experimental and 3 technical replicates. Data shown as mean±SD normalized to the respective conditions with maximum fluorescence (Tau^ΔK280^+14-3-3ζ or PKA-Tau^ΔK280^). **b,** FRAP of “maturing” Tau (left) or PKA-Tau (right; 10 μM; Tau with 2% Tau-Dylight488, middle panel) condensates formed in 25 mM HEPES, 1 mM DTT, pH 7.4, 5% PEG and with or without 14-3-3ζ (0 or 5 μM; with 2% 14-3-3ζ-DyLight650). Condensates were analyzed at 2 h and 6 h after formation. N=15-18 condensates per condition. Data shown as mean±SEM. **c,** FRAP of Tau variants (10 μM Tau with 2% Tau-Dylight488) and 14-3-3ζ (5 μM 14-3-3ζ with 2% 14-3-3ζ-DyLight650). N=15-18 condensates per condition. Data shown as mean±SEM. **d,** Representative images of condensates formed from fluorescently labeled Tau variants and 14-3-3ζ right before and after photobleaching and after 40 sec of recovery. Condensates formed in 25 mM HEPES, 1 mM DTT, pH 7.4 in the presence of 5% PEG. Scale bars = 5 μm.

Next, we investigated whether 14-3-3 interactions would alter Tau condensate maturation. The mobility of proteins within condensates is determined by the amount and strength of their interactions and for Tau typically decreases during condensate maturation. We used fluorescence recovery after photobleaching (FRAP) to evaluate the mobility (diffusion) and mobile fraction of Tau in and 14-3-3ζ in 14-3-3:Tau condensates. By bleaching small regions (<20% of condensates area), we particularly probed these parameters inside condensates. Both unphosphorylated Tau and 14-3-3ζ were highly mobile in condensates (mobile fractions: Tau 61%, t_1/2_=4s; 14-3-3ζ 85%, t_1/2_=7s) (Fig 6b,c). Similarly, phosphorylated non-binding mutant PKA-Tau^S214A/S324A^ demonstrated a comparable fraction of mobile PKA-Tau^S214A/S324A^ and a slightly reduced fraction of mobile 14-3-3ζ molecules (mobile fractions: PKA-Tau^S214A/S324A^ 64%, t_1/2_=7s; 14-3-3ζ 72%, t_1/2_=13s). By contrast, the mobile fractions in PKA-Tau:14-3-3ζ condensates was decreased (mobile fractions: PKA-Tau 50%, t_1/2_=6s; 14-3-3ζ 50%, t_1/2_=17s). This indicated that 14-3-3:Tau binding reduced the mobility of both binding partners in condensates. Notably, PKA-Tau^S214A/S324A^ recovered more slowly than non-phosphorylated Tau, suggesting that Tau phosphorylation (aside from S214 and S324) generally increased interactions in Tau condensates. These interactions were likely of electrostatic nature, given that PKA-Tau^S214A/S324A^ condensates were more sensitive to NaCl levels in the buffer than non-phosphorylated Tau condensates (Fig. 5h).

Lastly, to assess whether 14-3-3 binding would alter condensate maturation, we compared the time-dependent loss of Tau molecular mobility inside Tau and PKA-Tau condensates (measured by FRAP) in the absence and presence of 14-3-3ζ. Between 2 h and 6 h after condensate formation, Tau FRAP decreased similarly in all conditions tested (Fig. 6d), showing that neither the presence nor the binding of 14-3-3ζ changed Tau condensate maturation. This suggested that the progressive loss of Tau mobility in condensates depended on Tau:Tau and not Tau:14-3-3 interactions, and that Tau:14-3-3ζ condensates could still catalyze Tau aggregation.

## Discussion

### Regulation of Tau MT binding via 14-3-3 binding competition

Binding and stabilization of axonal MTs are the canonical functions of Tau^80,81^, derived from the high affinity of Tau to MTs both *in vitro* and in cells. Tau phosphorylation in the microtubule binding regions, which is largely comprised of TauRD, is generally assumed to be sufficient for Tau dissociation from MTs^19^. Our data suggest that, *in vitro* and in neurons, phospho-Tau can remain bound to MTs as long as it is not phosphorylated at main 14-3-3 binding sites, pS214 and pS324, in the presence of 14-3-3. Phosphorylation at these sites increases the affinity of phospho-Tau to 14-3-3, thereby allowing 14-3-3 to “scavenge” Tau_pS214/pS324_ from the MT surface. The exchange of Tau molecules between MTs and 14-3-3 appears to be tightly regulated, given the opposition in binding affinity; non-phosphorylated Tau has a high affinity to MTs (K_D_ < 1 μM)^9,67^ but does not bind 14-3-3ζ (Fig. 2), while the MT affinity of PKA-Tau is decreased (K_D_ ≍ 10 μM^68^) but its 14-3-3 affinity is increased (K_D_ ≍ 1 μM; Fig. 2). This interplay would limit the availability of unbound, aggregation-prone phospho-Tau in the cytosol and also ensure that phospho-Tau can efficiently leave the MT surface, for example, to allow for physiological MT dynamics (Fig. 7). These findings support our hypothesis that Tau-MT binding appears to relay not only on Tau phosphorylation but also on the presence of phospho-Tau binding partners, like 14-3-3.

**Figure 7.**
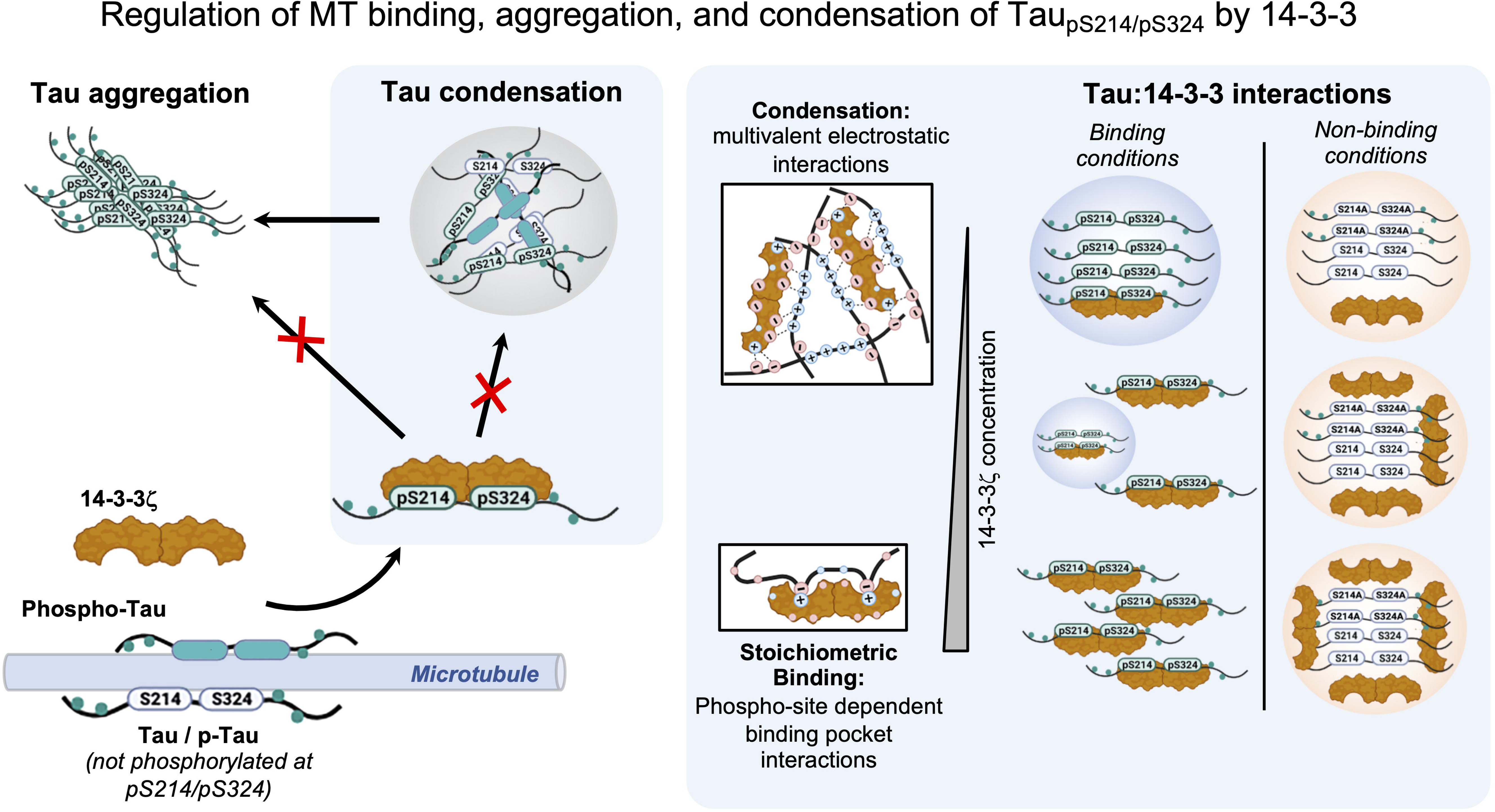
Regulation of Tau MT binding, aggregation and condensation by 14-3-3ζ. **Left:** 14-3-3ζ dimers form stable stoichiometric complexes with Tau pS214/pS324. This enables the efficient dissociation of Tau pS214/pS324 from MTs, which may contribute to ensure MT dynamics. In addition, the formation of 14-3-3ζ:Tau pS214/pS324 complexes prevents the aggregation of Tau in the cytosol by inhibiting phospho-Tau assembly into fibrillar aggregates and/or liquid-like condensates. In non-binding conditions, when Tau is not phosphorylated at S214 and S324, 14-3-3ζ co-condenses with Tau based on multivalent electrostatic interactions. These condensates attach to MTs, when present, and sequesters excess Tau from the solution. Whether 14-3-3ζ:Tau condensates have of cellular function remains to be clarified. **Right:** Interactions of Tau and 14-3-3 in stoichiometric binding vs. condensation conditions. In binding conditions, stoichiometric binding of Tau in the binding groove of 14-3-3ζ dimers precludes Tau condensation by depleting the pool of Tau molecules necessary to drive condensation. This is due to occupation of TauRD and the proline-rich region in Tau - domains that drive Tau condensation - by 14-3-3ζ. In non-binding-conditions, multivalent electrostatic interactions between the 14-3-3 surface and Tau drive co-condensation. Notably, 14-3-3 dimers in complex with Tau (or other clients) can also participate in co-condensation because their surface charges are still accessible.

It is important to be aware that Tau molecules can be phosphorylated at many sites and that each molecule may carry a differential pattern of ∼8 phosphorylations^21^ and/or other PTMs^16^, which may all influence Tau’s binding to other molecules, including different 14-3-3 isoforms and MTs. *Previous studies showed that PKA phosphorylation first introduces pS214, then rather simultaneously pS324, pS356, pS409, and pS416, and to a lower degree pS262 and other sites*^82^*. pS214 and pS324, and to a lesser degree pS356 (having a ∼4-fold lower individual binding affinity than the other two), are mainly relevant for 14-3-3 binding.* Excess phosphorylation by PKA was shown to weaken the affinity of individual Tau:14-3-3 interactions sites (e.g., pS324, pS356, pS409), but the affinities of pS214 and the *overall affinity between 14-3-3 and Tau seemed to be not much altered by higher PKA phosphorylation. Accordingly, we found that PKA phosphorylated phospho-null mutants lacking phosphorylation at pS214 and pS324 (Tau S214A/S324A) lose their ability to bind 14-3-3, however, they retain their MT binding, notably, while still being phosphorylated at other PKA target*.

*Additionally, the Tau:14-3-3/Tau:MT interaction balance is likely further tuned by other, non-PKA PTMs. Tau phosphorylation at common phosphorylation sites outside of the TauRD, introduced by CDK5 (mainly pS202, pT205, pS235, pS404*^83^*) and Fyn (pY18*^84^*), did not catalyze 14-3-3 binding, but were elsewhere reported to reduce MT binding*^13^. To which extend Tau PTMs other than pS214 and pS324 tune *the Tau:14-3-3/Tau:MT interaction balance – an thereby impact Tau aggregation -* needs further investigation. Additionally, Tau:14-3-3 interactions can also be tuned modifications on 14-3-3^82^.

One may also consider that excess of 14-3-3 could cause a depletion of Tau from the MT surface and, hence, a destabilization of the MT cytoskeleton. Similar observations have been made upon 14-3-3σ overexpression in cells and primary neurons^60^. Such effects of high 14-3-3 expression may have multiple reasons, given that 14-3-3 proteins are important hubs in cell biology that regulate the activity and availability of many proteins. For example, other studies showed that stabilizing 14-3-3 interactions using fusicoccin-A negatively affects the stress response regulator GCN1 and thereby improves axon outgrowth and regeneration in cultured neurons and mouse brain^85^. Tau phosphorylation is generally high in these conditions, i.e., in immature neurons^17^ with axon outgrowth and in injured neurons^86^, which may favor Tau binding to 14-3-3 proteins and other chaperoning proteins instead of MTs, thereby contributing to the MT dynamics necessary during axon outgrowth and repair.

### Inhibition of Tau condensation and aggregation by 14-3-3

When binding, 14-3-3ζ inhibits *in vitro* Tau amyloid aggregation (Fig. 6a;^49,51,87^) as well as condensation (Fig. 4;^76^). In contrast, 14-3-3 enhanced Tau aggregation under non-binding conditions and promoted Tau condensation at sub-stoichiometric 14-3-3 dimer concentrations and in non-binding conditions (i.e., non-phosphorylated Tau or PKA-Tau^S214A/S324A^). “Maturation” of Tau condensates and the accompanying risk of producing seeding-competent Tau species^3^ was not prevented by 14-3-3ζ. This suggests that, in order to prevent Tau condensation/aggregation, sufficient 14-3-3 molecules need to be present in the cytosol to complex Tau_pS214/pS324_. Accordingly, reduced 14-3-3 levels, increased (phospho-)Tau levels, and de-regulated Tau PTMs – as observed in AD^54^ – would increase the risk for uncontrolled cytosolic Tau condensation, condensate maturation, and eventually Tau seed formation.

To robustly prevent condensation and aggregation of Tau not binding 14-3-3, Tau needs to be chaperoned by other binding partners of similar or higher abundance and/or binding strength. In this regard, binding of Tau to MTs - its main cellular binding partner - may act as a sink for Tau that cannot be bound by 14-3-3, for example due to the lack of phosphorylation at S214 and S324, or by other chaperones.

### Regulation of Tau solubility via 14-3-3 client competition

Whether phosphorylated at S214 and S324 or not, the cellular pool of Tau available for condensation and aggregation can be influenced by the binding competition between 14-3-3 clients. Here, PKA-Tau and the pS2 Tau phospho-peptide seem to influence each other’s solubility (liquid-condensed vs. free soluble) based on their availability for 14-3-3 binding. Similar competition between Tau and many other competing 14-3-3 binding partners, including a number of condensing^74^ and aggregating^88–90^ proteins, could indirectly influence the condensation of each other. Cellular pro-pathological Tau condensation and aggregation could, thus, be tuned not only by its own phosphorylation at S214 and S324 but also by phosphorylation and availability of other 14-3-3 clients. Furthermore, since 14-3-3 bound to one client could co-condense with other clients, 14-3-3 proteins may generate phase separated catalytic hubs.

### Structural underpinnings of Tau binding to 14-3-3

Structure and binding mode of 14-3-3ζ dimers in complex with the pS2 peptide in this study were comparable to previous studies using other 14-3-3 isoforms and Tau constructs^22,44,60^. Extrapolation of the pS2:14-3-3 structure to full-length Tau_pS214/pS324_ indicates that 14-3-3 binding would cover and conformationally restrict large parts of the TauRD. This is lin ine with previous observations using NMR^82^. TauRD mediates Tau’s binding to MTs^14,91,92^, but it also comprises the hexapeptide amino acid motifs relevant for Tau aggregation^93–95^, builds the core of AD and FTD amyloid fibrils^64,65^, and is important for Tau condensation^72,96^. In addition, condensation of the intrinsically disordered protein Tau may require molecular flexibility and expansion^97^ that may be hindered upon binding to 14-3-3. It is important to note that the pS2 phospho-peptide used in this study consisted of Tau peptides around the two phospho-sites important for binding, connected by an artificial, short (11 aa, ∼3 nm) and unstructured linker. In full-length Tau, however, the same peptides are connected by ∼100 aa (also largely unstructured with some beta-structure^98^). This to some extend limits our interpretation of the binding affinity of pS2 to 14-3-3, which may bind more or less strongly than full-length Tau_pS214/pS324_. Thermal stability assays indicated that full-length PKA Tau may bind even stronger to 14-3-3ζ than the pS2 peptide. It is nevertheless clear that these two phospho-sites are important to the binding of 14-3-3ζ to Tau, and the interaction of 14-3-3ζ with the TauRD explains its compound impact on Tau MT binding, aggregation, and condensation.

### Relevance of 14-3-3 for Tau brain pathology

Tau protein aggregation is a pathological hallmark in >20 neurodegenerative brain diseases. Although numerous factors – mutations, PTMs, and polyanionic co-factors – can seemingly increase Tau aggregation, it is unclear what promotes the high solubility and prevents the aggregation of Tau in healthy neurons, notably, despite the rather high physiological concentration of Tau (∼2 μM; ^1–3^). Our data show that phospho-site dependent and specific, stoichiometric binding of 14-3-3 can keep Tau in solution by actively preventing its assembly into liquid-like condensates and amyloid-like fibrils (Fig. 6). This could explain why in the healthy brain, where most Tau molecules are somehow phosphorylated^99^, cytosolic phospho-Tau aggregation and condensation is efficiently prevented. In Alzheimer’s disease, where 14-3-3 proteins are reduced^54^ and Tau phosphorylation is generally increased, insufficient complexation of phospho-Tau by 14-3-3 could allow the accumulation of aggregation-capable free phospho-Tau in the cytosol. Increasing 14-3-3:Tau binding may contribute to prevent aberrant Tau aggregation as a therapeutic approach.

Our data show that fusicoccin, a molecule non-specifically increasing 14-3-3 interactions with its clients including Tau, increased levels of cytosolic soluble Tau and reduced the number of neurons developing Tau aggregates. Designing “molecular glues” that specifically stabilize 14-3-3:Tau interactions, facilitating Tau solubility even in situations of otherwise reduced protein homeostasis, could reduce Tau aggregation in the brain. In fact, similar approaches may also be explored for neurodegenerative conditions involving other condensing and aggregating client proteins of 14-3-3, for example, a-synuclein in Parkinson’s disease^58,88,100^. Notably, we could confirm the presence of the phospho-epitopes important for efficient 14-3-3 binding, Tau_pS214_ and Tau_pS324_, in human brain and cultured mouse neurons but not in other mammalian cell (Supplemental Fig. S5), suggesting a possible brain tissue specific role of 14-3-3 in promoting Tau solubility. Whether 14-3-3 promotes Tau solubility in other tissues, such as in kidney^101^ and testis^102,103^, needs to be tested.

In summary, our results propose a model in which 14-3-3 proteins make an important contribution to neuronal Tau solubility. Stoichiometric, phosphorylation-dependent binding of 14-3-3 enables dissociation of Tau_pS214/pS324_ from the MT surface, which may be important to enable MT dynamics. At the same time, the binding of 14-3-3 ensures the solubility of Tau_pS214/pS324_ in the cytosol by suppressing its condensation and amyloid aggregation. The seemingly-less “handing-over” of Tau between the MT surface and chaperoning 14-3-3 dimers could underlie pathological Tau aggregation occurring when Tau phosphorylation increases and 14-3-3 levels and other complementary homeostatic mechanisms start to deteriorate, for example in the AD brain.

## Methods

### Primary neurons expressing eGFP-Tau^P301L/S320F^

Primary hippocampal mouse neurons were prepared from dissected hippocampi of postnatal (P0-1) wild-type mice and grown under standard conditions in PDL-coated 8-well imaging dishes (ibidi). On DIV5, neurons were AAV transduced for the expression of EGFP-tagged mutant Tau (pAAV.CAG.EGFP-Tau^P301L/S320F^). Neurons were treated with BV02 (10 µM or 40 µM; #SML0140, Sigma) or fusicoccin (50 µM; # sc-200754, Santa Cruz) on DIV8 for 48 h. On DIV10, neurons were fixed in 4% PFA in PBS, counterstained with DAPI for 10 min, and imaged on a Nikon scanning Confocal A1Rsi+ or a Leica Stellaris Falcon microscope. A 10x air objective was use to acquire images of a large field of view (entire well of culture dish) for quantification, and a 60x oil objective to collect images of individual neurons.

### Immunofluorescence

Primary hippocampal mouse neurons (DIV 10) were washed with PBS before fixation with 4% PFA in PBS for 15 min, followed by a wash with TBS for 10 min. After fixation, cells were permeabilized using 0.5% Triton in PBS for 20 min and subsequently blocked with 3% Normal Goat Serum (NGS) in PBS for 1 h at room temperature. Incubation with the primary antibodies (mouse anti-Tau pSer202/pSer205 (AT8), Invitrogen # MN1020 (1:500)); rabbit anti-Tau_pS214_, Abcam #ab170892 (1:1000); rabbit anti-Tau_pS324_, Abcam #ab109401 (1:1000); chicken anti-Map2, Abcam #ab92434 (1:1000); rabbit anti-14-3-3ζ, Abcam #ab51129 (1:1000)) in 3% NGS in PBS was performed overnight at 4°C. After three washes in PBS for 10 min, neurons were incubated with secondary antibodies (DyLight405 goat anti-mouse IgG, Invitrogen #35501BID; AlexaFluor647 goat anti-chicken IgY, Invitrogen #A21449; AlexaFluor555 goat anti-rabbit IgG, Invitrogen # A27039 (all 1:2000)) in 3% NGS in PBS for 2 h at room temperature. After three washes for 10 min in PBS, nuclei were counterstained with DAPI (1:1000 in PBS) for 10Cmin. Cells were imaged on a Nikon scanning Confocal A1Rsi+ or a Leica Stellaris Falcon 8 microscope with a 60x oil objective. Images were processed and quantified with ImageJ.

### Co-immunoprecipitation

Primary cortical mouse neurons (DIV 9) were harvested from 6-well plates in lysis buffer (10 mM Tris/Cl pH7.5, 150 nM NaCl, 0.5 mM EDTA, 0.5 % NP40, phosphatase and protease inhibitors). Co-immunoprecipitation was performed using magnetic ProteinG Dynabeads (LifeTechnologies, #10003D) following the manufacturer protocol. In short, 30 µl bead slurry was washed with PBS + Tween (0.02%), then incubated for 30 min at room temperature under rotation with 8 μg of one of the following antibodies: rabbit anti-total Tau (DAKO A0024); rabbit anti-IgG isotype control (Millipore 12-370). Excess antibody in the supernatant was removed by pulling out the beads using a magnet before cell lysates (250 µg protein in 500 µl lysis buffer) were added to antibody-bound ProteinG Dynabeads overnight at 4°C under rotation. Magnetic beads were separated from supernatant (flow-through) using a magnet and beads were washed 3x with lysis buffer. Elution of proteins bound to beads was performed by resuspending the beads in 35 µl 2xSDS sample buffer followed by boiling at 95°C for 5 min. Samples without beads were separated by SDS-PAGE and WB 14-3-3 detected by Western Blot (mouse anti-pan 14-3-3 (H-8), Santa Cruz, #sc-1657 (1:800)).

### Protein purification

Recombinant human full-length Tau (2N4R, hTau40) and Tau^ΔK280^ were expressed in *E. coli* BL21 Star (DE3) (Invitrogen) cells as previously described^104^ and purified following an established protocol^55^. 14-3-3 protein zeta (14-3-3ζ) and 14-3-3ζ truncated after T234 (14-3-3ζΔC, for crystallography) were expressed in NiCo21 (DE3) competent cells and purified as previously described following an established protocol^105^. All recombinant purified and further modified proteins were stored aliquoted at −80°C.

### In vitro phosphorylation of Tau

Tau protein (6-8Cmg/ml) was incubated in phosphorylation buffer (25CmM HEPES, 100CmM NaCl, 5Cmg MgCl, 2CmM EGTA, 1CmM DTT, Protease Inhibitors) with recombinant PKA kinase (2500 U/mg, NEB-P600S) and 1CmM ATP overnight at 30°C and 250Crpm. To denature and remove the kinase from the sample, NaCl was added to a final concentration of 500 mM. The protein solution was boiled for 10 min at 95 °C, and spun at 100000 g for 40 min. The phosphorylated Tau in the supernatant was dialyzed against phosphate buffered saline (PBS) containing 1 mM DTT, or against 25CmM HEPES, 10mM NaCl, 1CmM DTT, pH 7.4 for LLPS assays.

### Western Blot

Tau variants (0.5 μg protein), cell and brain lysates (10 μg protein) were separated by SDS-PAGE (NuPage 4-12% Bis-Tris, Invitrogen) and blotted onto nitrocellulose membranes. The membrane was blocked with 3% BSA in PBS containing 0.05% Tween (PBS-T) at room temperature for 1 h, followed by the incubation with primary anti-phospho-Tau antibodies (rabbit anti-Tau_pS214_, Abcam, #ab170892 (1:1000); rabbit anti-Tau_pS324_, Abcam #ab109401 (1:5000), mouse anti-Tau_pS202/pT205_ (AT8), Invitrogen #MN1020 (1:1000)) or and anti-total Tau antibody (rabbit anti-human Tau, Dako #A0024 (1:5,000)) diluted in 3% BSA in PBS-T overnight at 4°C. After washing with PBS-T, membranes were incubated with HRP-conjugated secondary antibody (goat anti-rabbit IRDye680 and goat anti-mouse IRDye800, LI-COR) in 3% BSA in PBS-T for 1-2 h at room temperature, washed in PBS-T, and then imaged on a LI-COR ODYSSEY.

### Tau peptides

Acetylated and fluorescein (FITC)-labeled Tau peptides for crystallography and fluorescence anisotropy were purchased from GenScript (sequences: mono-phospho Tau pS214: SRTP{pSER}LPTPPTRE; mono-phospho Tau pS324: VTSKCG{pSER}LGNIHHK; bi-phospho pS2 peptide: SRTP{pSER}LPTPPTREGGGSGGGSGGGVTSKCG{pSER}LGNIHHK).

### Fluorescence anisotropy assay (FA)

14-3-3ζ was titrated in a 2-fold dilution series (starting at 400 μM 14-3-3ζ) to 10 nM of fluorescein labeled peptide (Tau_pS214_, Tau_pS324_, Tau_pS214/pS324_ (=pS2)) in FA buffer (10 mM HEPES pH 7.4, 150 mM NaCl, 50 µM TCEP, 0.1% (v/v) Tween20, 0.1% (w/v) BSA). Dilution series were made in polystyrene low-volume 384-well plates (Corning #4514, Black Round Bottom). Measurements were performed directly after plate preparation, using a Tecan SPARK plate reader at room temperature (λ_ex_: 485±20 nm; λ_em_: 535±25 nm; mirror: automatic; flashes: 30; settle time: 1 ms; gain: optimal; Z-position: calculated from well). Wells containing only fluorescein labelled peptide were used to set as G-factor calibrated from these wells. All data were analyzed using GraphPad Prism (10.0.1) and fitted with a four-parameter logistic model (4PL) to determine binding affinities (dissociation constant, KD). All results are based on three independent experiments from which the average and standard deviations for each KD were determined using excel. Tau competition assays were performed in a similar way but using full-length Tau constructs (WT, PKA-WT, PKA-Tau^S214A^, PKA-Tau^S324A^, PKA-Tau^S214A/S324A^) titrated in a 2-fold dilution series (starting from 50 μM) to 1 μM 14-3-3ζ and 10 nM fluorescein labelled Tau^pS214/pS324^ peptide.

### Differential Scanning Fluorimetry (DSF)

DSF was performed using 40 μl samples containing 2.5 µM 14-3-3ζ and 25 μM Tau and 5x ProteoOrange (Lumiprobe, 5000x stock in DMSO) in 10 mM HEPES, 150 mM NaCl, 50 μM TCEP (pH 7.4). The samples were heated from 35 °C to 79°C at a rate of 0.3°C per 15 s in a CFX96 Touch Real-Time PCR Detection System (Bio-Rad). Fluorescence intensity was determined using excitation (λ_em_=525/20 nm) and emission (λ_ex_=570/20 nm) filters. Based on these melting curves, the negative derivative melting curve is obtained from which the melting temperature Tm was determined. All described melting temperatures are based on three independent experiments, from which the average and standard deviations were determined.

### Microtubule pelleting assay

To assess the binding of Tau to MTs, we used a MT binding protein spin-down assay kit (# BK029, Cytoskeleton, Inc.) according to the manufacturer’s instructions. Briefly, MTs were assembled from soluble tubulin in MT assembly buffer (80 mM PIPES pH 7.0, 2 mM MgCl2, 0.5 mM EGTA) at 35°C for 20 min and stabilized with Taxol, yielding a concentration of approximately 5×10^11^ MT/ml. For MT binding, 0.4 μg recombinant Tau (Tau or PKA-Tau) with or without 1 μg 14-3-3ζ were incubated with 20 μl preformed MTs for 30 min. Prepared reactions (50 μl) were carefully added on top 100 μl cushion buffer (80 mM PIPES pH 7.0, 1 mM MgCl2, 1 mM EGTA, 60% glycerol) in small ultracentrifuge tubes (#343775, Beckmann) and centrifuged at 100000 g at room temperature for 40 min. After centrifugation, supernatant (50 μl) and pellet were separated and SDS sample buffer was added. Samples were analyzed by SDS-PAGE and western blot using total human Tau (#835201, Biolegend), 14-3-3 (#ab51129, Abcam), and α-tubulin (#T6074, Sigma) antibodies.

### In vitro microtubule bundle formation

Bovine tubulin (5 μM total tubulin, 10% Alexa594-labeled tubulin (PUR-032005 and PUR-059405, PurSolutions):) and 1 mM GTP in freshly prepared and filtered polymerization buffer (BRB80, 1 mM DTT, pH 6.8) were added to Tau (25 μM; 2% DyLight488-Tau) with and without 14-3-3ζ (12.5 μM; 2% DyLight650-14-3-3ζ) and in the presence of 5% (w/v) PEG8000. After 45 min, samples were pipetted into multi-well glass bottom dishes (μ-Slide Angiogenesis glass bottom dishes, Ibidi) and MT bundles, Tau, and 14-3-3ζ were imaged using a confocal spinning disc microscope (CSU-X, Nikon) with a 60x oil objective.

### 14-3-3ζ X-ray crystallography data collection and refinement binary structure

The 14-3-3ζΔC protein and the acetylated Tau^pS214/pS324^ peptide were dissolved in complexation buffer (20 mM HEPES pH 7.5, 2 mM MgCl2 and 100 μM TCEP) and mixed in a 1:1 molar stoichiometry (protein:peptide) at a final protein concentration of 15 mg/ml. The complex was set up for sitting-drop crystallization and crystals were grown within one month at 4°C in 0.2M Sodium Fluoride, 0.1M bis-tris propane pH 6.5, 20% w/v PEG 3350. Crystals were fished and flash-cooled in liquid nitrogen. X-ray diffraction (XRD) data was collected at the European Synchrotron Radiation Facility (ESRF) beamline ID23-1, Grenoble, France. autoPROC software (version 1.1.7) was used to index and integrate the diffraction data^106^. The data was further processed using the CCP4i2 suite (version 8.0.002)^107^. Scaling was performed using AIMLESS^108,109^. MolRep^110,111^ was used for phasing, using PDB ID 5D2D as template. REFMAC (version 5) ^112,113^ was used for initial structure refinement. Correct peptide sequences were modelled in the electron density in COOT (version 0.9.8.1)^114^. Alternating cycles of model improvement and refinement were performed using COOT and REFMAC. Figures were generated with PyMOL (version 2.5.2). 2Fo-Fc electron density maps were contoured at 1σ. See Table S and for XRD data collection, structure determination, and refinement statistics. The structures were submitted to the PDB with IDs: 8QDV.

### Fluorescent labeling of Tau and 14-3-3ζ

Fluorescent labelling of Tau and 14-3-3ζ protein for microscopy assays was done using amine-reactive DyLight405 or DyLight488-NHS ester (Thermo Scientific) following the manufacturer instructions and the established protocol^55^.

### Condensate formation and fluorescence microscopy

Prior to condensation assays, all proteins used were dialyzed against 25 mM HEPES, 10 mM NaCl, 1 mM DTT, pH7.4 and stored at -80°C. LLPS of Tau (10 μM, 2% DyLight488-Tau) with 14-3-3ζ (1-20 μM, 2% DyLight650-14-3-3z) variants was performed in 25 mM HEPES, 1 mM DTT, pH7.4 buffer (final NaCl concentration in sample 1-5 mM) in the presence of 5 % (w/vol) PEG-8000, except otherwise indicated. Directly after LLPS induction, 2-3 μl sample were pipetted onto glass-bottom dishes for imaging (TC-treated Miltenyi CG 1.5, diameter = 3.5 mm). Imaging dishes were immediately closed and equipped with a ddH_2_O-soaked paper tissue lining the inner edges to avoid evaporation of the LLPS sample. Imaging was performed 1 h after droplet formation either on a widefield epifluorescence microscope (Ti2, Nikon) or a spinning disk confocal microscope (CSU-X, Nikon) equipped with 40x air or 60x oil objective, respectively.

### Fluorescence recovery after photobleaching (FRAP)

Condensates (containing 2%Tau-DyLight488 and 2%14-3-3ζ-DyLight650) were imaged in a triggered acquisition mode before and directly after bleaching for 40 sec (round ROIs, cross sections 1-2 μM) with a 650 nm laser, followed by a 488 nm laser (90% intensity; 3 loops). In each field of view, same-sized background ROI (outside condensates) and non-bleached reference-ROI (inside different condensates) were measured. All FRAP measurements were background corrected and normalized to the background corrected reference signal. FRAP experiments were performed on a spinning disk confocal microscope (CSU-X, Nikon) using a 60x oil objective.

### Thioflavin-T Tau aggregation assay

Tau (10CμM Tau^ΔK280^ and PKA-Tau^ΔK280^) aggregation in the presence and absence of 14-3-3ζ (20 μM) was induced by adding heparin (0.115 mg/ml, MWC=C8–25 kDa (∼7 μM), Applichem) in PBS, pH7.4, 1 mM DTT. Thioflavin-T (ThioT, 50 μM, Sigma) was added to detect the aggregation. Samples were prepared in triplicates, pipetted into black clear-bottom 384-well μClear plates (Greiner), and ThioT fluorescence (λ_Ex_C=C440Cnm, λ_Em_C=C485Cnm) was recorded in a plate reader (Infinite M Plex, Tecan) following a 5 sec shake every 15 min at 37°C.

## Data availability

All raw data (images, FRAP data, WBs) are available upon request. The X-ray structure of 14-3-3ζ in complex with pS2 peptide is available via the PDB; 8QDV.

## Acknowledgements

We thank Michelle Arkin (UCSF) for stimulating discussions. The crystallography data collection was performed at the European Synchrotron Radiation Facility (ESRF), Grenoble, France. We would like to thank Lindsay McGregor for assistance in using beamline ID23-1. Beamtime was allocated for proposal MX-2407 (doi 10.15151/ESRF-ES-1006014569). Microscopy was performed at the AMBIO imaging core of the Charité University Berlin, and at the imaging core of the MDC-BIMSB (Leica Stellaris Falcon setup). AAVs were produced at the viral core facility (VCF) of the Charité University Berlin.

## Author Contributions

J.H., performed *in vitro* LLPS and aggregation, FRAP, and MT imaging experiments, drafted initial Figures and helped writing the manuscript. M.C.M.v.d.O, determined crystal structure of 14-3-3ζ in complex with pS2, performed fluorescence anisotropy and thermal stability measurements, helped writing the manuscript. L.D., performed neuron experiments and MT pelleting assay, and helped reviewing the manuscript. L.R. performed *in vitro* condensation assays and edited the manuscript. L.L, helped with LLPS experiments and designed 14-3-3ζ mutant. R.P-L. and G.N., helped with neuron experiments. E.S., cloned and generated the mutant Tau AAV. S.M. helped with data analysis and reviewing the manuscript. C.O., helped supervising experiments, provided funding, reviewed manuscript. L.B., designed study, provided funding, supervised team at University Eindhoven, reviewed manuscript. S.W., designed study, provided funding, supervised team at DZNE Berlin, wrote and reviewed manuscript.

## Supplemental Figure legends

**Supplemental Figure S1. Tau phosphorylation and TaupS214/pS324 and pS2 peptide binding to 14-3-3**ζ.

**a,** Western blots of cell lysates from cultured primary mouse neurons show phosphorylation of endogenous mouse Tau at S214 and S324. **b,** Amino acid sequences (letter code) of Tau phospho-peptides. Phospho-sites are marked in red letters, linker region in blue letters. **c,** Independent experimental replicates of fluorescence anisotropy measurements for FITC-labeled Tau peptides pS2, Tau_pS214_, and Tau_pS324_ (related to Fig. 2e). Data shown as mean±SD, N=3 technical replicates per experiment. **d,** Independent experimental replicates of fluorescence anisotropy measurements for full-length PKA-Tau variants competing with pS2-FITC bound to 14-3-3ζ. Data shown as mean±SD, N=3 technical replicates per experiment. **e,** Phos-Tag gel shows efficient phosphorylation of Tau by PKA and Cdk5 compared to non-phosphorylated Tau.

**Supplemental Figure S2. 14-3-3**ζ **alone does not condensate.**

Representative images of different 14-3-3ζ concentrations (incl. 2% 14-3-3ζ-DyLight650) in condensation assay buffer (HEPES, pH7.4, 5% PEG). 14-3-3ζ and 14-3-3ζ^R127A^ alone do not form condensates even at 100 μM. Scale bars = 20 μm.

**Supplemental Figure S3.**

**a,** Second 14-3-3ζ dimer part of the asymmetric unit in complex with Tau phospho-peptide pS2. Cartoon plot with semi-transparent surface of the 14-3-3ζ dimer (gray) complexed with the pS214 and pS324 binding sites of the pS2 (pink rods). **b+c,** Close-up of the 14-3-3ζ binding groove in complex with individual pS2 binding sites (pink rods). Final 2Fo-Fc electron density map of pS2 is shown as blue mesh (contoured at 1s). **d,** Top view of 14-3-3ζ (grey semi-transparent surface) in complex with pS2 binding motifs around pS214 and pS324 (pink rods) connected by the unstructured linker (purple dotted line). **e,** Crystal packing of pS2:14-3-3ζ complexes. Two 14-3-3ζ dimers forming a tetramer (gray semi-transparent surfaces) are bound to two two pS2 peptides (green and pink rods).

**Supplemental Figure S4. 14-3-3**ζ **induces Tau condensation at net charge matching.** Representative images of 10 μM non-phosphorylated Tau with increasing 14-3-3ζ concentrations (0, 1, 5, 10, 20 μM in 25 mM HEPES, pH 7.4, 1 mM DTT, 5% (w/v) PEG; 2% Tau-DyLight488, 2% 14-3-3ζ-DyLight647). Scale bars = 20 μm.

**Supplemental Figure S5. Tau pS214/pS324 in cell and brain lysates.**

**a+b,** Western blots of cell lysates from mouse hippocampal primary neurons (**a**) and different mammalian cell lines and AD brain (**b**) confirm the presence of both Taup_S214_ and Tau_pS324_ phosphorylation only in neurons and human brain.

**Supplemental Table 1. Data collection and refinement statistics of the crystal structure.** Statistics for the highest-resolution shell are shown in parentheses.

