## Supplemental Material for "Stoichiometric 14-3-3ζ binding promotes phospho-Tau microtubule dissociation and reduces aggregation and condensation"

### Supplemental Figure S1

**a** Co-IP of 14-3-3 with Tau from neuron lysates

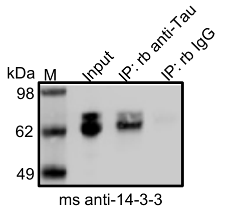


**b** Tau phospho-epitopes modified by kinaes in vitro

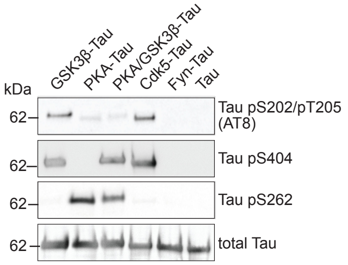


**c** Tau phospho-peptide amino acid sequences

| | |
|---|---|
| Tau_pS214_pS324 (**pS2**) | SRTP(pS)LPTPPTREGGGSGGGSGGGVTSKCG(pS)LGNIHHK |
| Tau_pS214 | SRTP(pS)LPTPPTRE |
| Tau_pS324 | VTSKCG(pS)LGNIHHK |

**d** Tau phospho-peptide binding assay (replicates)

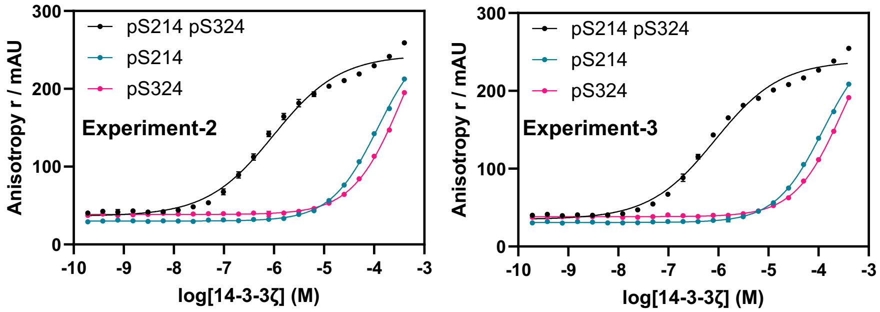


**e** pS2 peptide competition assay (replicates)

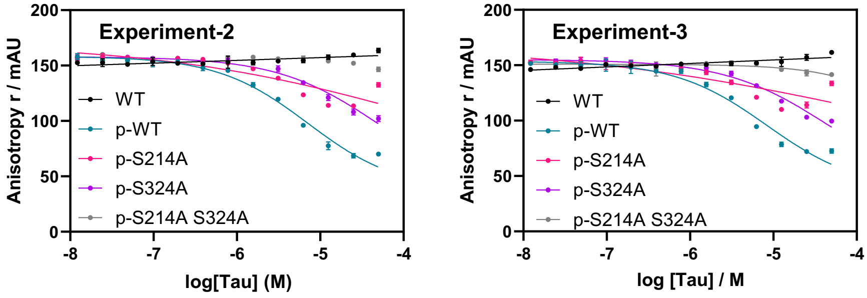


**f** Phos-Tag gel

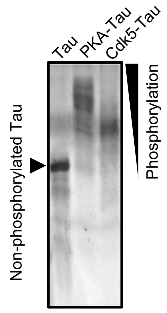

### Supplemental Figure S2

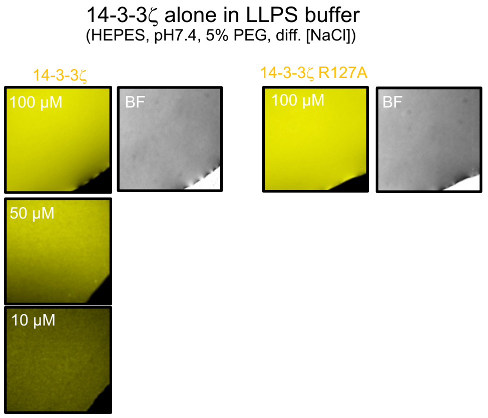

### Supplemental Figure S3

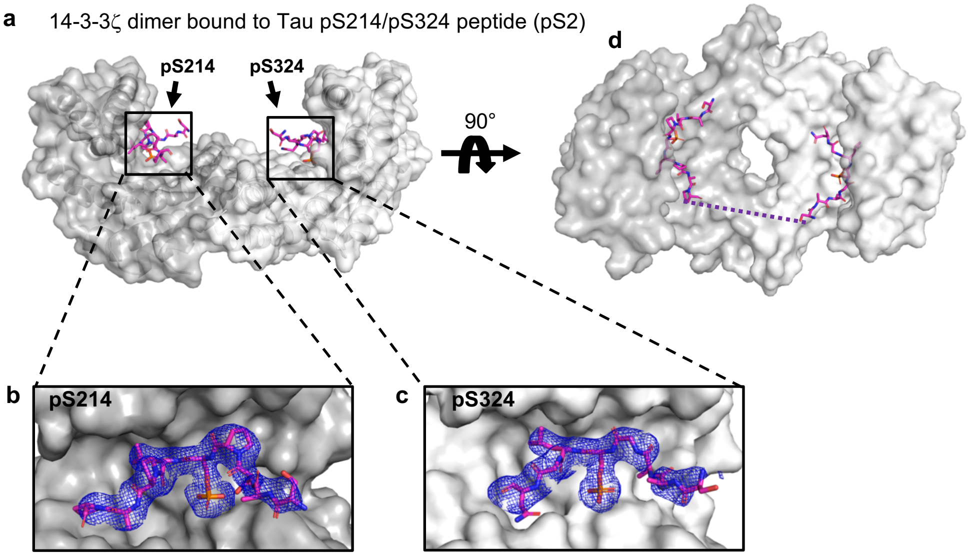


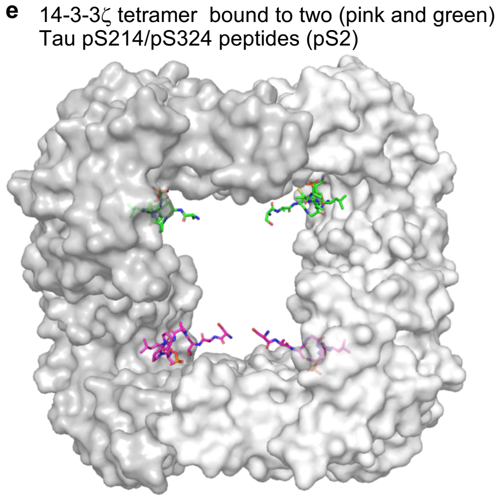

### Supplemental Figure S4

Condensation of non-phosphorylated Tau

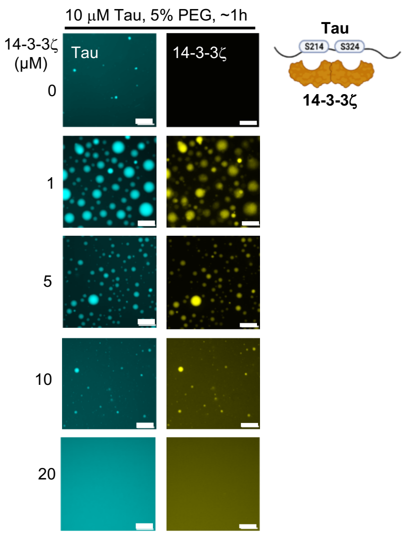

### Supplemental Figure S5

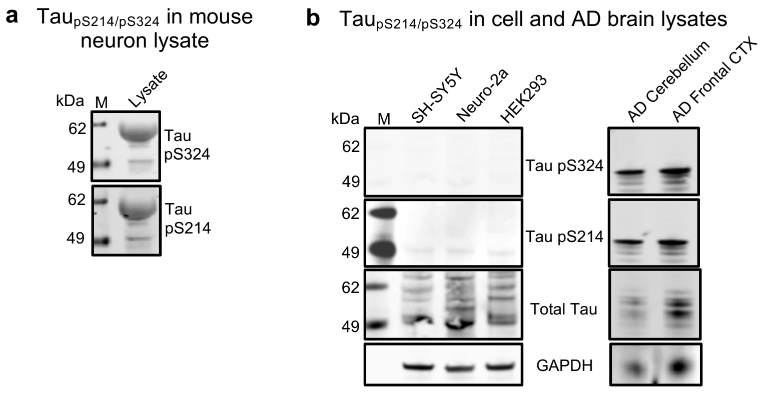
